## Supplementary Information I for "Engineered cell differentiation and sexual reproduction in probiotic and mating yeasts"

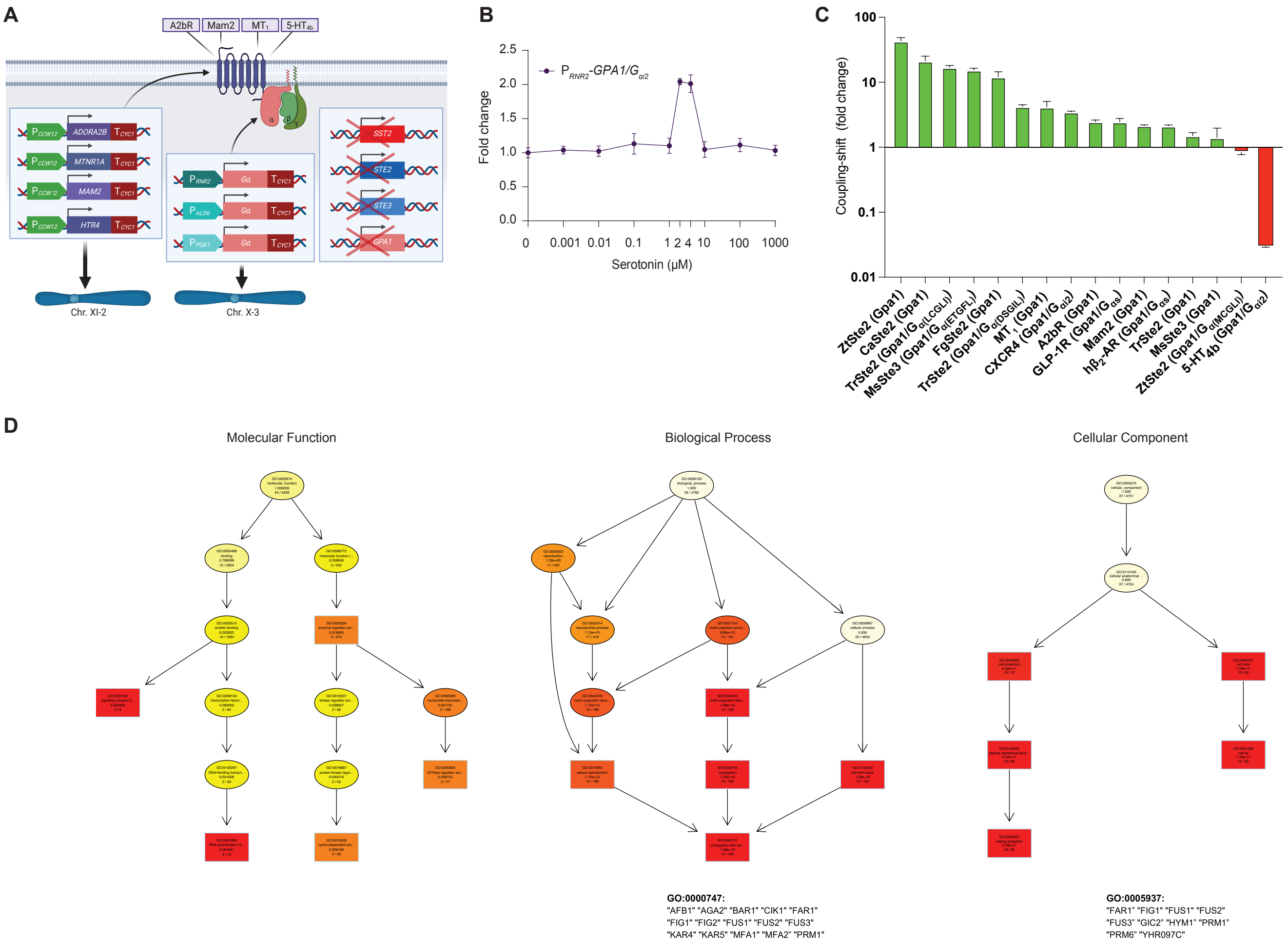


**Supplementary Figure S1. A.** Schematic overview on engineering of biosensing yeast strains. Heterologous GPCRs (hGPCRs) are overexpressed from the *CCW12* promoter and integrated in the genome, and the *CYC1* terminator attenuated transcription in all designs. Genome-integrated $G_\alpha$ subunits were expressed from the *RNR2, ALD6, or PGK1* promoters. Four native genes, *SST2*, *STE2*, *STE3*, and *GPA1*, were genetically deleted, and $G_\beta$ and $G_\gamma$ subunits are illustrated in a trimeric complex with $G_\alpha$ subunit. **B.** Fold change from 0-1000 µM serotonin supplementation in strain CPK165. **C.** Coupling-shifts are presented as log-scaled fold changes in fluorescence from a $P_{FUS1}$-*GFP* reporter following hGPCR integration (+hGPCR: strains SBY143, SBY146, CPK153, CPK156, CPK159, CPK165, and CPK450-459) over background (no hGPCR: SBY123, CPK131, CPK134, CPK343, CPK347, CPK350, and CPK424). Specific $G_\alpha$ subunits are indicated for each hGPCR presented in the plot. Means and standard deviations represent three biological replicates. **D.** The GO trees induced by the top 5 GO terms for “molecular function”, “biological process”, and “cellular component” respectively. Rectangles indicate the 5 most significant terms. The color represents the relative significance, ranging from dark red (most significant) to bright yellow (least significant). For each node, the first two lines show the GO information. The third line is the raw p-value, and the fourth line shows the number of significant genes and the total number of genes annotated to the respective GO term.

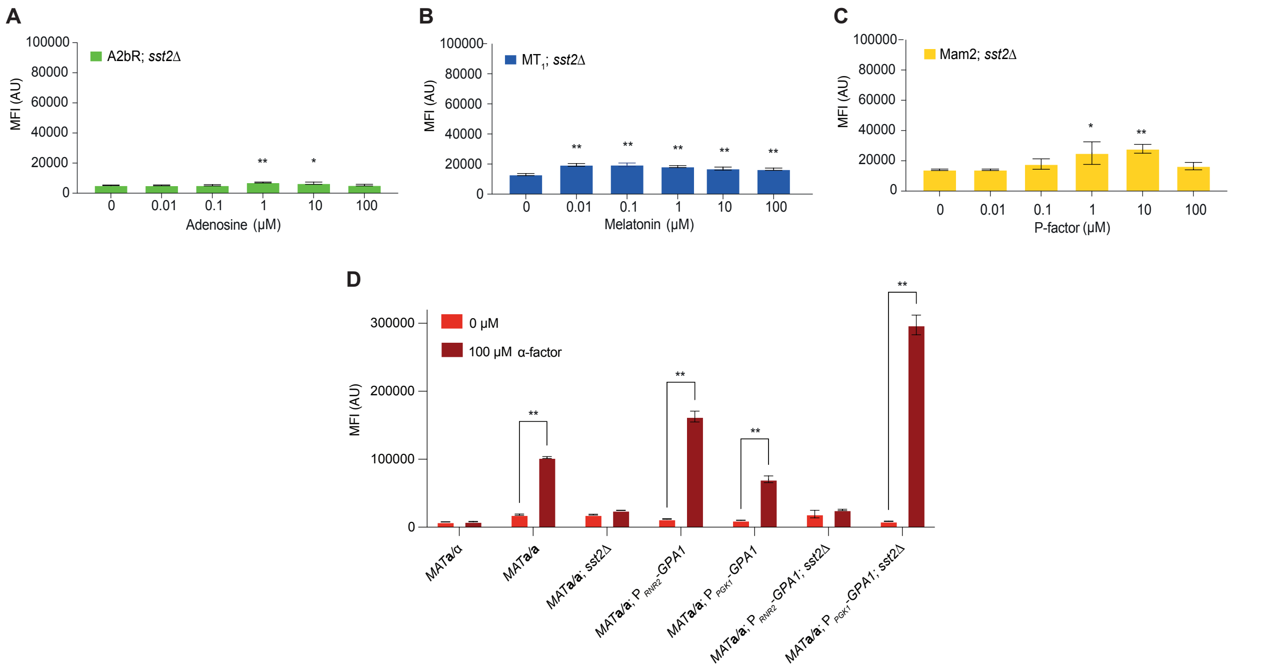


**Supplementary Figure S2. A-C:** Median fluorescence intensity (MFI) shown in artificial units (AU) from plasmid-based $P_{FUS1}$-*GFP* reporter expression in *S. boulardii sst2Δ* biosensing strains for **A.** A2bR sensing adenosine (SB45), **B.** $MT_1$ sensing melatonin (SB46), and **C.** Mam2 sensing P-factor (SB47). **D.** Mating pathway stimulation with α-factor in engineered *S. boulardii* strains: SB14 (*MAT***a**/α), SB17 (*MAT***a**/**a**), SB36 (*MAT***a**/**a***; sst2Δ*), SB40 (*MAT***a**/**a**; $P_{RNR2}$-*GPA1*), SB39 (*MAT***a**/**a**; $P_{PGK1}$-*GPA1*), SB38 (*MAT***a**/**a**; $P_{RNR2}$-*GPA1; sst2Δ*), and SB37 (*MAT***a**/**a**; $P_{PGK1}$-*GPA1; sst2Δ*). Means and standard deviations represent three biological replicates. Statistical significance was determined relative to no ligand supplementation (0 μM) controls using one-way or two-way analysis of variance (ANOVA) for (**A-C**) and for (**D**), respectively, using GraphPad Prism (*p > 0.05, **p ≤ 0.01).

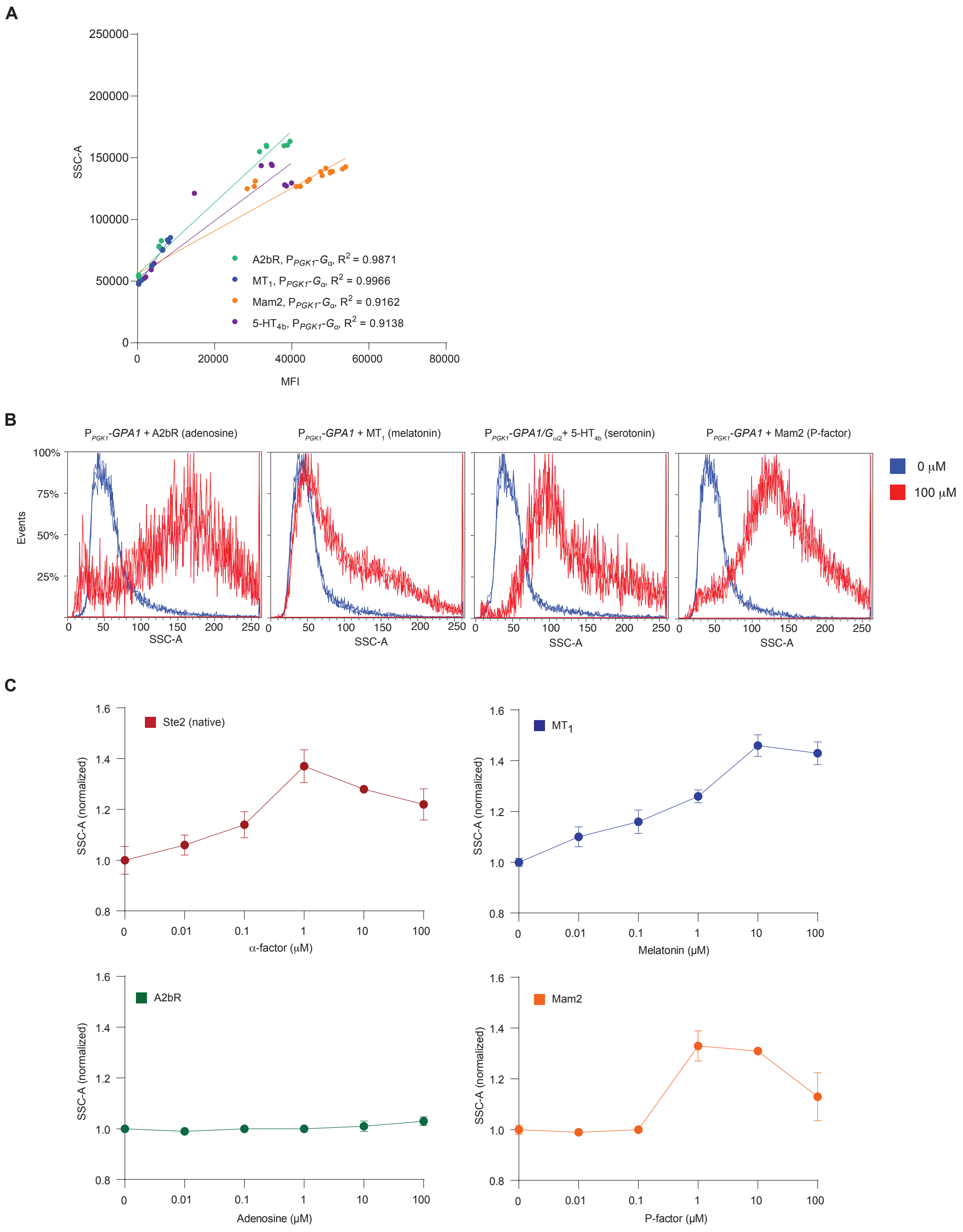


**Supplementary Figure S3. A.** Side-scatter (SSC-A) as a linear function of median fluorescence intensity (MFI) for strains CPK155, CPK158, CPK161, and CPK167. **B.** Histograms showing increased SSC-A during hGPCR-signaling in strains CPK155, CPK158, CPK161, and CPK167 from incubation with cognate ligands (100 µM, red) as compared to no hGPCR-signaling (0 µM, blue). **C.** SSC-A changes with cognate ligand concentration in biosensing *S. boulardii* (SB17 and SB48-50). SSC-A is normalized internally for each biosensing strain to no ligand supplementation controls (0 µM).

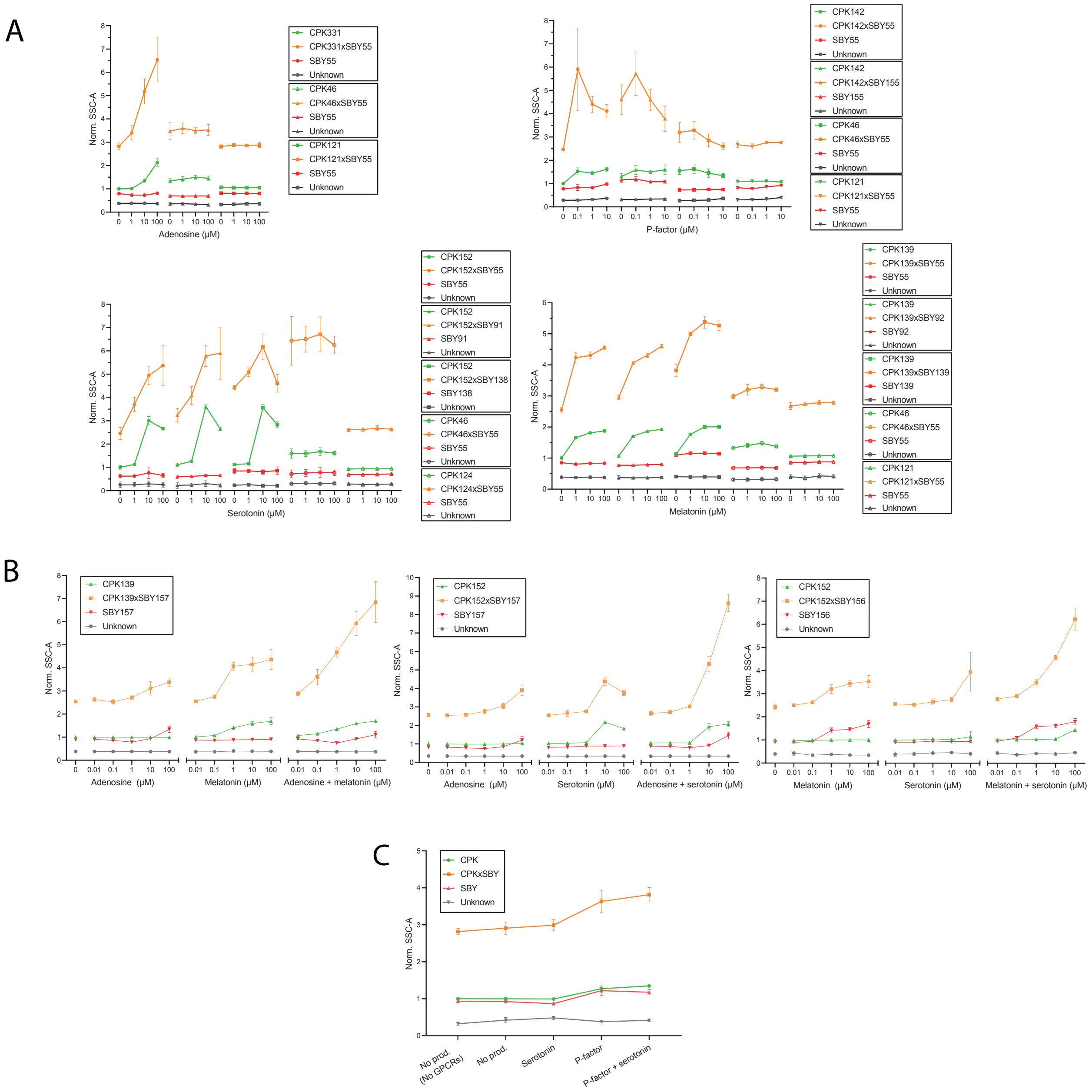


**Supplementary Figure S4.** Alterations in cellular morphology during synthetic mating in the formation of diploids (CPKxSBY) and shmooing haploids (CPK and SBY), interpreted by side-scatter area (SSC-A). SSC-A was measured simultaneously with mating data acquisition. Cells are identified by flow cytometry (see Methods). SSC-A is normalized to the examined CPK-strain without supplementation or production of ligand. CPK46xSBY55 and CPK121xSBY55 or CPK124xSBY55 were used as references for positive and negative mating pair controls with relevant ligand supplementation, respectively. Results are presented as means with standard deviations determined from five biological replicates. **A**. Normalized SSC-A for semi-synthetic mating trials for adenosine in Fig. 4B, P-factor in Fig. 4C, melatonin in Fig. 4D, and serotonin in Fig. 4E. **B**. Normalized SSC-A for full synthetic mating trials for adenosine and melatonin in Fig. 5B, adenosine and serotonin in Fig. 5C, and melatonin and serotonin in Fig. 5D. **C**. Normalized SSC-A for full autonomous mating trials in Fig. 5E.

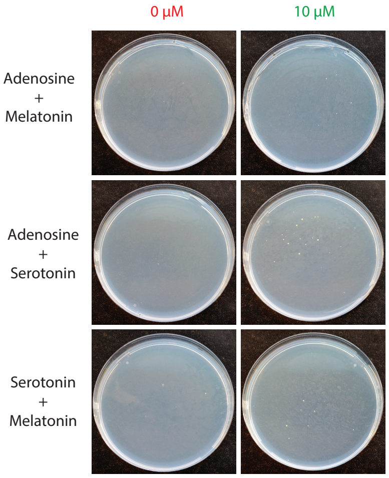


**Supplementary Figure S5.** Representative pictures of plated full synthetic mating trials for supplementation (10 µM) of both ligands in combination, or no supplementation (0 µM), for crosses with strains corresponding to CPK139xSBY157 (adenosine+melatonin), CPK152xSBY157 (adenosine+serotonin), and CPK152xSBY156 (melatonin+serotonin). CPK139 and CPK152 were transformed with plasmid pEDJ400 (*URA3*) prior to synthetic mating to allow for diploid selection on SC-UW. 95 µl co-culture was plated on each SC-UW plate, and pictures were taken following incubation at 30 °C for 7 days.

### Supplementary Table S4

| Strain | *Genotype* | Plasmid(s) | Parental strain | Reference |
|---|---|---|---|---|
| yWS677 | *MAT**a**; sst2Δ0; far1Δ0; bar1Δ0; ste2Δ0; ste12Δ0; gpa1Δ0; ste3Δ0; mf (alpha)1Δ0; mf(alpha)2Δ0; mfa1Δ0; mfa2Δ0; gpr1Δ0; gpa2Δ0* | N/A | N/A | Shaw et al. 2019 |
| CEN.PK2-1C | *MAT**a**; his3D1; leu2-3_112; ura3-52; trp1-289; MAL2-8c; SUC2* | N/A | N/A | EUROSCARF |
| BY4741 | *MAT**a;** his3Δ1; leu2Δ0; met15Δ0; ura3Δ0* | N/A | N/A | EUROSCARF |
| SBY1 | *MAT**a**; his3Δ1; leu2Δ0; met15Δ0; ura3Δ0* | pEDJ391 | BY4741 | This study |
| SBY2 | *MAT**a**; his3Δ1; leu2Δ0; met15Δ0; ura3Δ0; sst2Δ* | pEDJ391 | SBY1 | This study |
| SBY3 | *MAT**a**; his3Δ1; leu2Δ0; met15Δ0; ura3Δ0; sst2Δ; ste3Δ* | pEDJ391 | SBY2 | This study |
| SBY4 | *MAT**a**; his3Δ1; leu2Δ0; met15Δ0; ura3Δ0; sst2Δ; ste2Δ; ste3Δ* | pEDJ391 | SBY3 | This study |
| SBY16 | *MAT**a**; his3Δ1; leu2Δ0; met15Δ0; ura3Δ0; sst2Δ; ste2Δ; ste3Δ; fus1::GFP* | pEDJ391 | SBY4 | This study |
| SBY53 | *MAT**a**; his3Δ1; leu2Δ0; met15Δ0; ura3Δ0; pTEF1-mRuby2-tADH1-XII-2* | pEDJ391 | SBY1 | This study |

| | | | | |
|---|---|---|---|---|
| SBY55 | *MATα; his3Δ1; leu2Δ0; met15Δ0; ura3Δ0; pTEF1-mRuby2-tADH1-XII-2* | pEDJ391 | SBY53 | This study |
| SBY91 | *MATα; his3Δ1; leu2Δ0; met15Δ0; ura3Δ0 ; pTEF1-mRuby2-tADH1-XII-2; tADH1-HsDDC<-pTDH3-pTEF1->SmTPH-tCYC1-XI-3; tADH1-RnPTS<-pTEF1-PGK1p->RnSPR-tCYC1-X-4; tADH1-PaPCBD1<-pTEF1-PGK1p->RnDHPR-tCYC1-XII-4* | N/A | SBY55 | This study |
| SBY92 | *MATα; his3Δ1; leu2Δ0; met15Δ0; ura3Δ0; pTEF1-mRuby2-tADH1-XII-2; tADH1-HsDDC<-pTDH3-pTEF1->SmTPH-tCYC1-XI-3; tADH1-RnPTS<-pTEF1-PGK1p->RnSPR-tCYC1-X-4; tADH1-PaPCBD1<-pTEF1-PGK1p->RnDHPR-tCYC1-XII-4; tADH1-HsASMT<-pTEF1-PGK1p->BtAANAT-tCYC1-X-2* | N/A | SBY91 | This study |
| SBY104 | *MAT**a**; his3Δ1; leu2Δ0; met15Δ0; ura3Δ0; sst2Δ; ste2Δ; ste3Δ; pPGK1-GPA1-tCYC1-X-3* | pEDJ391 | SBY4 | This study |
| SBY108 | *MAT**a**; his3Δ1; leu2Δ0; met15Δ0; ura3Δ0; sst2Δ; ste2Δ; ste3Δ; fus1:: GFP; pRNR2-GPA1-tCYC1-X-3* | pEDJ391 | SBY16 | This study |
| SBY116 | *MAT**a**; his3Δ1; leu2Δ0; met15Δ0; ura3Δ0; sst2Δ; ste2Δ; ste3Δ; pPGK1-GPA1-tCYC1-X-3; gpa1Δ0* | pEDJ391 | SBY104 | This study |

| | | | | |
|---|---|---|---|---|
| SBY120 | *MAT**a**; his3Δ1; leu2Δ0; met15Δ0; ura3Δ0; sst2Δ; ste2Δ; ste3Δ; fus1::GFP; pRNR2-GPA1-tCYC1-X-3; gpa1Δ0* | pEDJ391 | SBY108 | This study |
| SBY123 | *MAT**a**; his3Δ1; leu2Δ0; met15Δ0; ura3Δ0; sst2Δ; ste2Δ; ste3Δ; fus1::GFP; pRNR2-GPA1/Gαs-tCYC1-X-3; gpa1Δ0* | pEDJ391 | SBY120 | This study |
| SBY128 | *MAT**a**; his3Δ1; leu2Δ0; met15Δ0; ura3Δ0; sst2Δ; ste2Δ; ste3Δ; pPGK1-GPA1-tCYC1-X-3; gpa1Δ0; pTEF1-mKATE2-tCYC1-XII-2* | pEDJ391 | SBY116 | This study |
| SBY138 | *MATα; his3Δ1; leu2Δ0; met15Δ0; ura3Δ0 ; pTEF1-mRuby2-tADH1-XII-2; tADH1-HsDDC<-pTDH3-pTEF1->SmTPH-tCYC1-XI-3; tADH1-RnPTS<-pTEF1-PGK1p->RnSPR-tCYC1-X-4; tADH1-PaPCBD1<-pTEF1-PGK1p->RnDHPR-tCYC1-XII-4; Ty2-LoxP-KlURA3-TAG-pPGK1-SmTPH-tCYC1* | N/A | SBY91 | This study |
| SBY139 | *MATα; his3Δ1; leu2Δ0; met15Δ0; ura3Δ0; pTEF1-mRuby2-tADH1-XII-2; tADH1-HsDDC<-pTDH3-pTEF1->SmTPH-tCYC1-XI-3; tADH1-RnPTS<-pTEF1-PGK1p->RnSPR-tCYC1-X-4; tADH1-PaPCBD1<-pTEF1-PGK1p->RnDHPR-tCYC1-XII-4; tADH1-HsASMT<-pTEF1-PGK1p->BtAANAT-tCYC1-X-2; Ty2-LoxP-KlURA3-TAG-pPGK1-SmTPH-tCYC1* | N/A | SBY92 | This study |

| | | | | |
|---|---|---|---|---|
| SBY143 | *MAT**a**; his3Δ1; leu2Δ0; met15Δ0; ura3Δ0; sst2Δ; ste2Δ; ste3Δ; fus1::GFP; pRNR2-GPA1/Gαs-tCYC1-X-3; gpa1Δ0; pCCW12:GLP-1R:tCYC1-XI-2* | N/A | SBY123 | This study |
| SBY146 | *MAT**a**; his3Δ1; leu2Δ0; met15Δ0; ura3Δ0; sst2Δ; ste2Δ; ste3Δ; fus1::GFP; pRNR2-GPA1/Gαs-tCYC1-X-3; gpa1Δ0; pCCW12:ADRB2:tCYC1-XI-2* | N/A | SBY123 | This study |
| SBY155 | *MATα; his3Δ1; leu2Δ0; met15Δ0; ura3Δ0 ; pTEF1ro-mRuby2-XII-2; pTEF1-α-leader_P-factor-tCYC1* | N/A | SBY55 | This study |
| SBY156 | *MATα; his3Δ1; leu2Δ0; met15Δ0; ura3Δ0; sst2Δ; ste2Δ; ste3Δ; X-3-pPGK1:GPA1:tCYC1; gpa1Δ0 ; pTEF1-mKATE2-tCYC1-XII-2; pCCW12-MTNR1A-tCYC1-XI-2* | N/A | SBY128 | This study |
| SBY157 | *MATα; his3Δ1; leu2Δ0; met15Δ0; ura3Δ0; sst2Δ; ste2Δ; ste3Δ; X-3-pPGK1:GPA1:tCYC1; gpa1Δ0 ; pTEF1-mKATE2-tCYC1-XII-2; pCCW12-ADORA2B-tCYC1-XI-2* | N/A | SBY128 | This study |
| SBY172 | *MATα; his3Δ1; leu2Δ0; met15Δ0; ura3Δ0; sst2Δ; ste2Δ; ste3Δ; pPGK1-GPA1-tCYC1-X-3; gpa1Δ0 ; pTEF1-mKATE2-tCYC1-XII-2* | pEDJ391 | SBY128 | This study |

| | | | | |
|---|---|---|---|---|
| SBY173 | *MATα; his3Δ1; leu2Δ0; met15Δ0; ura3Δ0; sst2Δ; ste2Δ; ste3Δ; pPGK1-GPA1-tCYC1-X-3; gpa1Δ0 ; pTEF1-mKATE2-tCYC1-XII-2; pCCW12-MAM2-tCYC1-XI-2* | N/A | SBY172 | This study |
| SBY174 | *MATα; his3Δ1; leu2Δ0; met15Δ0; ura3Δ0; sst2Δ; ste2Δ; ste3Δ; pPGK1-GPA1-tCYC1-X-3; gpa1Δ0 ; pTEF1-mKATE2-tCYC1-XII-2; pCCW12-MAM2-tCYC1-XI-2; tADH1-HsDDC<-pTDH3-pTEF1->SmTPH-tCYC1-XI-3; tADH1-RnPTS<-pTEF1-PGK1p->RnSPR-tCYC1-X-4; tADH1-PaPCBD1<-pTEF1-PGK1p->RnDHPR-tCYC1-XII-4* | N/A | SBY173 | This study |
| SBY175 | *MATα; his3Δ1; leu2Δ0; met15Δ0; ura3Δ0; sst2Δ; ste2Δ; ste3Δ; pPGK1-GPA1-tCYC1-X-3; gpa1Δ0 ; pTEF1-mKATE2-tCYC1-XII-2; pCCW12-MAM2-tCYC1-XI-2; tADH1-HsDDC<-pTDH3-pTEF1->SmTPH-tCYC1-XI-3; tADH1-RnPTS<-pTEF1-PGK1p->RnSPR-tCYC1-X-4; tADH1-PaPCBD1<-pTEF1-PGK1p->RnDHPR-tCYC1-XII-4; Ty2-LoxP-KlURA3-TAG-pPGK1-SmTPH-tCYC1* | N/A | SBY174 | This study |
| CPK1 | *MAT**a**; his3D1; leu2-3_112; ura3-52; trp1-289; MAL2-8c; SUC2* | pEDJ391 | CEN.PK2-1C | This study |
| CPK2 | *MAT**a**; his3D1; leu2-3_112; ura3-52; trp1-289; MAL2-8c; SUC2; sst2Δ* | pEDJ391 | CPK1 | This study |

| | | | | |
|---|---|---|---|---|
| CPK3 | *MAT**a**; his3D1; leu2-3_112; ura3-52; trp1-289; MAL2-8c; SUC2; sst2Δ; ste3Δ* | pEDJ391 | CPK2 | This study |
| CPK4 | *MAT**a**; his3D1; leu2-3_112; ura3-52; trp1-289; MAL2-8c; SUC2; sst2Δ; ste2Δ; ste3Δ* | pEDJ391 | CPK3 | This study |
| CPK16 | *MAT**a**; his3D1; leu2-3_112; ura3-52; trp1-289; MAL2-8c; SUC2; sst2Δ; ste2Δ; ste3Δ; fus1::GFP* | pEDJ391 | CPK4 | This study |
| CPK46 | *MAT**a**; his3D1; leu2-3_112; ura3-52; trp1-289; MAL2-8c; SUC2; pTEF1-GFP-tCYC1-XII-5* | N/A | CPK1 | This study |
| CPK88 | *MAT**a**; his3D1; leu2-3_112; ura3-52; trp1-289; MAL2-8c; SUC2; ste3Δ0; ste2Δ0; sst2Δ0* | pEDJ391 | CEN.PK2-1C | This study |
| CPK109 | *MAT**a**; his3D1; leu2-3_112; ura3-52; trp1-289; MAL2-8c; SUC2; ste3Δ0; ste2Δ0; sst2Δ0; pPGK1-GPA1-tCYC1-X-3* | pEDJ391 | CPK88 | This study |
| CPK112 | *MAT**a**; his3D1; leu2-3_112; ura3-52; trp1-289; MAL2-8c; SUC2; ste3Δ0; ste2Δ0; sst2Δ0; pPGK1-GPA1/Gαi2-tCYC1-X-3* | pEDJ391 | CPK88 | This study |

| | | | | |
|---|---|---|---|---|
| CPK115 | *MAT**a**; his3D1; leu2-3_112; ura3-52; trp1-289; MAL2-8c; SUC2; ste3Δ0; ste2Δ0; sst2Δ0; pPGK1-GPA1-tCYC1-X-3; gpa1Δ0* | pEDJ391 | CPK109 | This study |
| CPK118 | *MAT**a**; his3D1; leu2-3_112; ura3-52; trp1-289; MAL2-8c; SUC2; ste3Δ0; ste2Δ0; sst2Δ0; pPGK1-GPA1/Gαi2-tCYC1-X-3; gpa1Δ0* | pEDJ391 | CPK112 | This study |
| CPK119 | *MAT**a**; his3D1; leu2-3_112; ura3-52; trp1-289; MAL2-8c; SUC2; ste3Δ0; ste2Δ0; sst2Δ0; pRNR2-GPA1-tCYC1-X-3; gpa1Δ0; pTEF1-GFP-tCYC1-XII-5* | pEDJ391 | CPK113 | This study |
| CPK121 | *MAT**a**; his3D1; leu2-3_112; ura3-52; trp1-289; MAL2-8c; SUC2; ste3Δ0; ste2Δ0; sst2Δ0; pPGK1-GPA1-tCYC1-X-3; gpa1Δ0; pTEF1-GFP-tCYC1-XII-5* | pEDJ391 | CPK115 | This study |
| CPK124 | *MAT**a**; his3D1; leu2-3_112; ura3-52; trp1-289; MAL2-8c; SUC2; ste3Δ0; ste2Δ0; sst2Δ0; pPGK1-GPA1/Gαi2-tCYC1-X-3; gpa1Δ0; pTEF1-GFP-tCYC1-XII-5* | pEDJ391 | CPK118 | This study |
| CPK125 | *MAT**a**; his3D1; leu2-3_112; ura3-52; trp1-289; MAL2-8c; SUC2; sst2Δ; ste2Δ; ste3Δ; fus1::GFP; pRNR2-GPA1-tCYC1-X-3* | pEDJ391 | CPK16 | This study |

| | | | | |
|---|---|---|---|---|
| CPK126 | *MAT**a**; his3D1; leu2-3_112; ura3-52; trp1-289; MAL2-8c; SUC2; sst2Δ; ste2Δ; ste3Δ; fus1::GFP; pALD6-GPA1-tCYC1-X-3* | pEDJ391 | CPK16 | This study |
| CPK127 | *MAT**a**; his3D1; leu2-3_112; ura3-52; trp1-289; MAL2-8c; SUC2; sst2Δ; ste2Δ; ste3Δ; fus1::GFP; pPGK1-GPA1-tCYC1-X-3* | pEDJ391 | CPK16 | This study |
| CPK128 | *MAT**a**; his3D1; leu2-3_112; ura3-52; trp1-289; MAL2-8c; SUC2; sst2Δ; ste2Δ; ste3Δ; fus1::GFP; pRNR2-GPA1/Gαi2-tCYC1-X-3* | pEDJ391 | CPK16 | This study |
| CPK129 | *MAT**a**; his3D1; leu2-3_112; ura3-52; trp1-289; MAL2-8c; SUC2; sst2Δ; ste2Δ; ste3Δ; fus1::GFP; pALD6-GPA1/Gαi2-tCYC1-X-3* | pEDJ391 | CPK16 | This study |
| CPK130 | *MAT**a**; his3D1; leu2-3_112; ura3-52; trp1-289; MAL2-8c; SUC2; sst2Δ; ste2Δ; ste3Δ; fus1::GFP; pPGK1-GPA1/Gαi2-tCYC1-X-3* | pEDJ391 | CPK16 | This study |
| CPK131 | *MAT**a**; his3D1; leu2-3_112; ura3-52; trp1-289; MAL2-8c; SUC2; sst2Δ; ste2Δ; ste3Δ; fus1::GFP; pRNR2-GPA1-tCYC1-X-3; gpa1Δ0* | pEDJ391 | CPK125 | This study |

| | | | | |
|---|---|---|---|---|
| CPK132 | *MAT**a**; his3D1; leu2-3_112; ura3-52; trp1-289; MAL2-8c; SUC2; sst2Δ; ste2Δ; ste3Δ; fus1::GFP; pALD6-GPA1-tCYC1-X-3; gpa1Δ0* | pEDJ391 | CPK126 | This study |
| CPK133 | *MAT**a**; his3D1; leu2-3_112; ura3-52; trp1-289; MAL2-8c; SUC2; sst2Δ; ste2Δ; ste3Δ; fus1::GFP; pPGK1-GPA1-tCYC1-X-3; gpa1Δ0* | pEDJ391 | CPK127 | This study |
| CPK134 | *MAT**a**; his3D1; leu2-3_112; ura3-52; trp1-289; MAL2-8c; SUC2; sst2Δ; ste2Δ; ste3Δ; fus1::GFP; pRNR2-GPA1/Gαi2-tCYC1-X-3; gpa1Δ0* | pEDJ391 | CPK128 | This study |
| CPK135 | *MAT**a**; his3D1; leu2-3_112; ura3-52; trp1-289; MAL2-8c; SUC2; sst2Δ; ste2Δ; ste3Δ; fus1::GFP; pALD6-GPA1/Gαi2-tCYC1-X-3; gpa1Δ0* | pEDJ391 | CPK129 | This study |
| CPK136 | *MAT**a**; his3D1; leu2-3_112; ura3-52; trp1-289; MAL2-8c; SUC2; sst2Δ; ste2Δ; ste3Δ; fus1::GFP; pPGK1-GPA1/Gαi2-tCYC1-X-3; gpa1Δ0* | pEDJ391 | CPK130 | This study |
| CPK139 | *MAT**a**; his3D1; leu2-3_112; ura3-52; trp1-289; MAL2-8c; SUC2; ste3Δ0; ste2Δ0; sst2Δ0; pPGK1-GPA1-tCYC1-X-3; gpa1Δ0; pTEF1-GFP-tCYC1-XII-5; pCCW12-MTNR1A-tCYC1-XI-2* | N/A | CPK121 | This study |

| | | | | |
|---|---|---|---|---|
| CPK142 | *MAT**a**; his3D1; leu2-3_112; ura3-52; trp1-289; MAL2-8c; SUC2; ste3Δ0; ste2Δ0; sst2Δ0; pPGK1-GPA1-tCYC1-X-3; gpa1Δ0; pTEF1-GFP-tCYC1-XII-5; pCCW12-MAM2-tCYC1-XI-2* | N/A | CPK121 | This study |
| CPK152 | *MAT**a**; his3D1; leu2-3_112; ura3-52; trp1-289; MAL2-8c; SUC2; ste3Δ0; ste2Δ0; sst2Δ0; pPGK1-GPA1/Gαi2-tCYC1-X-3; gpa1Δ0; pTEF1-GFP-tCYC1-XII-5; pCCW12-HTR4-tCYC1-XI-2* | N/A | CPK124 | This study |
| CPK153 | *MAT**a**; his3D1; leu2-3_112; ura3-52; trp1-289; MAL2-8c; SUC2; sst2Δ; ste2Δ; ste3Δ; fus1::GFP; pRNR2-GPA1-tCYC1-X-3; gpa1Δ0; pCCW12-MTNR1A-tCYC1-XI-2* | N/A | CPK131 | This study |
| CPK154 | *MAT**a**; his3D1; leu2-3_112; ura3-52; trp1-289; MAL2-8c; SUC2; sst2Δ; ste2Δ; ste3Δ; fus1::GFP; pALD6-GPA1-tCYC1-X-3; gpa1Δ0; pCCW12-MTNR1A-tCYC1-XI-2* | N/A | CPK132 | This study |
| CPK155 | *MAT**a**; his3D1; leu2-3_112; ura3-52; trp1-289; MAL2-8c; SUC2; sst2Δ; ste2Δ; ste3Δ; fus1::GFP; pPGK1-GPA1-tCYC1-X-3; gpa1Δ0; pCCW12-MTNR1A-tCYC1-XI-2* | N/A | CPK133 | This study |

| | | | | |
|---|---|---|---|---|
| CPK156 | *MAT**a**; his3D1; leu2-3_112; ura3-52; trp1-289; MAL2-8c; SUC2; sst2Δ; ste2Δ; ste3Δ; fus1::GFP; pRNR2-GPA1-tCYC1-X-3; gpa1Δ0; pCCW12-MAM2-tCYC1-XI-2* | N/A | CPK131 | This study |
| CPK157 | *MAT**a**; his3D1; leu2-3_112; ura3-52; trp1-289; MAL2-8c; SUC2; sst2Δ; ste2Δ; ste3Δ; fus1::GFP; pALD6-GPA1-tCYC1-X-3; gpa1Δ0; pCCW12-MAM2-tCYC1-XI-2* | N/A | CPK132 | This study |
| CPK158 | *MAT**a**; his3D1; leu2-3_112; ura3-52; trp1-289; MAL2-8c; SUC2; sst2Δ; ste2Δ; ste3Δ; fus1::GFP; pPGK1-GPA1-tCYC1-X-3; gpa1Δ0; pCCW12-MAM2-tCYC1-XI-2* | N/A | CPK133 | This study |
| CPK159 | *MAT**a**; his3D1; leu2-3_112; ura3-52; trp1-289; MAL2-8c; SUC2; sst2Δ; ste2Δ; ste3Δ; fus1::GFP; pRNR2-GPA1-tCYC1-X-3; gpa1Δ0; pCCW12-ADORA2B-tCYC1-XI-2* | N/A | CPK131 | This study |
| CPK160 | *MAT**a**; his3D1; leu2-3_112; ura3-52; trp1-289; MAL2-8c; SUC2; sst2Δ; ste2Δ; ste3Δ; fus1::GFP; pALD6-GPA1-tCYC1-X-3; gpa1Δ0; pCCW12-ADORA2B-tCYC1-XI-2* | N/A | CPK132 | This study |

| | | | | |
|---|---|---|---|---|
| CPK161 | *MAT**a**; his3D1; leu2-3_112; ura3-52; trp1-289; MAL2-8c; SUC2; sst2Δ; ste2Δ; ste3Δ; fus1::GFP; pPGK1-GPA1-tCYC1-X-3; gpa1Δ0; pCCW12-ADORA2B-tCYC1-XI-2* | N/A | CPK133 | This study |
| CPK165 | *MAT**a**; his3D1; leu2-3_112; ura3-52; trp1-289; MAL2-8c; SUC2; sst2Δ; ste2Δ; ste3Δ; fus1::GFP; pRNR2-GPA1/Gαi2-tCYC1-X-3; gpa1Δ0; pCCW12-HTR4-tCYC1-XI-2* | N/A | CPK134 | This study |
| CPK166 | *MAT**a**; his3D1; leu2-3_112; ura3-52; trp1-289; MAL2-8c; SUC2; sst2Δ; ste2Δ; ste3Δ; fus1::GFP; pALD6-GPA1/Gαi2-tCYC1-X-3; gpa1Δ0; pCCW12-HTR4-tCYC1-XI-2* | N/A | CPK135 | This study |
| CPK167 | *MAT**a**; his3D1; leu2-3_112; ura3-52; trp1-289; MAL2-8c; SUC2; sst2Δ; ste2Δ; ste3Δ; fus1::GFP; pPGK1-GPA1/Gαi2-tCYC1-X-3; gpa1Δ0; pCCW12-HTR4-tCYC1-XI-2* | N/A | CPK136 | This study |
| CPK331 | *MAT**a**; his3D1; leu2-3_112; ura3-52; trp1-289; MAL2-8c; SUC2; ste3Δ0; ste2Δ0; sst2Δ0; pPGK1-GPA1-tCYC1-X-3; gpa1Δ0; pTEF1-GFP-tCYC1-XII-5; pCCW12-ADORA2B-tCYC1-XI-2* | N/A | CPK121 | This study |

| | | | | |
|---|---|---|---|---|
| CPK343 | *MAT**a**; his3D1; leu2-3_112; ura3-52; trp1-289; MAL2-8c; SUC2; sst2Δ; ste2Δ; ste3Δ; fus1::GFP; pRNR2-GPA1/Gα (LCGLI)-tCYC1-X-3; gpa1Δ0* | N/A | CPK131 | This study |
| CPK347 | *MAT**a**; his3D1; leu2-3_112; ura3-52; trp1-289; MAL2-8c; SUC2; sst2Δ; ste2Δ; ste3Δ; fus1::GFP; pRNR2-GPA1/Gα (DSGIL)-tCYC1-X-3; gpa1Δ0* | N/A | CPK131 | This study |
| CPK350 | *MAT**a**; his3D1; leu2-3_112; ura3-52; trp1-289; MAL2-8c; SUC2; sst2Δ; ste2Δ; ste3Δ; fus1::GFP; pRNR2-GPA1/Gα (ETGFL)-tCYC1-X-3; gpa1Δ0* | N/A | CPK131 | This study |
| CPK424 | *MAT**a**; his3D1; leu2-3_112; ura3-52; trp1-289; MAL2-8c; SUC2; sst2Δ; ste2Δ; ste3Δ; fus1::GFP; pRNR2-GPA1/Gα (MCGLI)-tCYC1-X-3; gpa1Δ0* | N/A | CPK131 | This study |
| CPK448 | *MATα; his3D1; leu2-3_112; ura3-52; trp1-289; MAL2-8c; SUC2; ste3Δ0; ste2Δ0; sst2Δ0; pPGK1-GPA1/Gαi2-tCYC1-X-3; gpa1Δ0; pTEF1-GFP-tCYC1-XII-5; pCCW12-HTR4-tCYC1-XI-2; pTEF1-α-leader_P-factor-tCYC1* | N/A | CPK152 | This study |
| CPK450 | *MAT**a**; his3D1; leu2-3_112; ura3-52; trp1-289; MAL2-8c; SUC2; sst2Δ; ste2Δ; ste3Δ; fus1::GFP; pRNR2-GPA1-tCYC1-X-3; gpa1Δ0; pCCW12-CaSTE2-tCYC1-XI-2* | N/A | CPK131 | This study |

| | | | | |
|---|---|---|---|---|
| CPK451 | *MAT**a**; his3D1; leu2-3_112; ura3-52; trp1-289; MAL2-8c; SUC2; sst2Δ; ste2Δ; ste3Δ; fus1::GFP; pRNR2-GPA1-tCYC1-X-3; gpa1Δ0; pCCW12-FgSTE2-tCYC1-XI-2* | N/A | CPK131 | This study |
| CPK452 | *MAT**a**; his3D1; leu2-3_112; ura3-52; trp1-289; MAL2-8c; SUC2; sst2Δ; ste2Δ; ste3Δ; fus1::GFP; pRNR2-GPA1-tCYC1-X-3; gpa1Δ0; pCCW12-ZtSTE2-tCYC1-XI-2* | N/A | CPK131 | This study |
| CPK453 | *MAT**a**; his3D1; leu2-3_112; ura3-52; trp1-289; MAL2-8c; SUC2; sst2Δ; ste2Δ; ste3Δ; fus1::GFP; pRNR2-GPA1-tCYC1-X-3; gpa1Δ0; pCCW12-TrSTE2-tCYC1-XI-2* | N/A | CPK131 | This study |
| CPK454 | *MAT**a**; his3D1; leu2-3_112; ura3-52; trp1-289; MAL2-8c; SUC2; sst2Δ; ste2Δ; ste3Δ; fus1::GFP; pRNR2-GPA1-tCYC1-X-3; gpa1Δ0; pCCW12-MsSTE3-tCYC1-XI-2* | N/A | CPK131 | This study |
| CPK455 | *MAT**a**; his3D1; leu2-3_112; ura3-52; trp1-289; MAL2-8c; SUC2; sst2Δ; ste2Δ; ste3Δ; fus1::GFP; pRNR2-GPA1/Gαi2-tCYC1-X-3; gpa1Δ0; pCCW12:CXCR4: tCYC1-XI-2* | N/A | CPK134 | This study |

| | | | | |
|---|---|---|---|---|
| CPK456 | *MAT**a**; his3D1; leu2-3_112; ura3-52; trp1-289; MAL2-8c; SUC2; sst2Δ; ste2Δ; ste3Δ; fus1::GFP; pRNR2-GPA1/Gα (MCGLI)-tCYC1-X-3; gpa1Δ0; pCCW12-ZtSTE2-tCYC1-XI-2* | N/A | CPK424 | This study |
| CPK457 | *MAT**a**; his3D1; leu2-3_112; ura3-52; trp1-289; MAL2-8c; SUC2; sst2Δ; ste2Δ; ste3Δ; fus1::GFP; pRNR2-GPA1/Gα (LCGLI)-tCYC1-X-3; gpa1Δ0; pCCW12-TrSTE2-tCYC1-XI-2* | N/A | CPK343 | This study |
| CPK458 | *MAT**a**; his3D1; leu2-3_112; ura3-52; trp1-289; MAL2-8c; SUC2; sst2Δ; ste2Δ; ste3Δ; fus1::GFP; pRNR2-GPA1/Gα (DSGIL)-tCYC1-X-3; gpa1Δ0; pCCW12-TrSTE2-tCYC1-XI-2* | N/A | CPK347 | This study |
| CPK459 | *MAT**a**; his3D1; leu2-3_112; ura3-52; trp1-289; MAL2-8c; SUC2; sst2Δ; ste2Δ; ste3Δ; fus1::GFP; pRNR2-GPA1/Gα (ETGFL)-tCYC1-X-3; gpa1Δ0; pCCW12-MsSTE3-tCYC1-XI-2* | N/A | CPK350 | This study |
| Sb.MYA-796 | *MAT**a**/α* | N/A | N/A | ATCC MYA-796 |
| DD277 | *MAT**a**/α; ura3Δ* | N/A | N/A | This study |
| DD313 | *MAT**a**/α; ura3Δ; his3Δ* | N/A | N/A | This study |
| SB4 | *MAT**a**/α; ura3Δ; his3Δ* | pDAM215 | DD313 | This study |

| | | | | |
|---|---|---|---|---|
| SB5 | *MAT**a**/α; ura3Δ; his3Δ; sst2Δ* | pDAM215 | SB4 | This study |
| SB6 | *MAT**a**/α; ura3Δ; his3Δ; sst2Δ; ste3Δ* | pDAM215 | SB5 | This study |
| SB8 | *MAT**a**/**a**; ura3Δ; his3Δ* | N/A | SB4 | This study |
| SB9 | *MAT**a**/**a**; ura3Δ; his3Δ; sst2Δ* | N/A | SB5 | This study |
| SB14 | *MAT**a**/α; ura3Δ; his3Δ* | pDAM194 | Sb.MYA-796 | This study |
| SB17 | *MAT**a**/**a**; ura3Δ; his3Δ* | pDAM194 | SB8 | This study |
| SB19 | *MAT**a**/α; ura3Δ; his3Δ; sst2Δ; ste3Δ; pPGK1-GPA1-tCYC1-X-3; gpa1Δ0* | pDAM215 | SB6 | This study |
| SB20 | *MAT**a**/α; ura3Δ; his3Δ; sst2Δ; ste3Δ; pRNR2-GPA1-tCYC1-X-3; gpa1Δ0* | pDAM215 | SB6 | This study |
| SB21 | *MAT**a**/α; ura3Δ; his3Δ; pPGK1-GPA1-tCYC1-X-3; gpa1Δ0* | pDAM215 | SB4 | This study |
| SB22 | *MAT**a**/α; ura3Δ; his3Δ; pRNR2-GPA1-tCYC1-X-3; gpa1Δ0* | pDAM215 | SB4 | This study |
| SB23 | *MAT**a**/α; ura3Δ; his3Δ; sst2Δ; ste3Δ; pPGK1-GPA1-tCYC1-X-3; gpa1Δ0; ste2Δ* | pDAM215 | SB19 | This study |
| SB24 | *MAT**a**/**a**; ura3Δ; his3Δ; sst2Δ; ste3Δ; pRNR2-GPA1-tCYC1-X-3; gpa1Δ0* | N/A | SB20 | This study |

| | | | | |
|---|---|---|---|---|
| SB25 | *MAT**a**/α; ura3Δ; his3Δ; pPGK1-GPA1-tCYC1-X-3; gpa1Δ0; ste2Δ* | pDAM215 | SB21 | This study |
| SB26 | *MAT**a**/**a**; ura3Δ; his3Δ; pRNR2-GPA1-tCYC1-X-3; gpa1Δ0* | N/A | SB22 | This study |
| SB30 | *MAT**a**/**a**; ura3Δ; his3Δ; sst2Δ; ste3Δ; pPGK1-GPA1-tCYC1-X-3; gpa1Δ0* | N/A | SB19 | This study |
| SB31 | *MAT**a**/**a**; ura3Δ; his3Δ; sst2Δ; ste3Δ; pPGK1-GPA1-tCYC1-X-3; gpa1Δ0; ste2Δ* | N/A | SB23 | This study |
| SB33 | *MAT**a**/**a**; ura3Δ; his3Δ; pPGK1-GPA1-tCYC1-X-3; gpa1Δ0; ste2Δ* | N/A | SB25 | This study |
| SB35 | *MAT**a**/**a**; ura3Δ; his3Δ; pPGK1-GPA1-tCYC1-X-3; gpa1Δ0* | N/A | SB21 | This study |
| SB36 | *MAT**a**/**a**; ura3Δ; his3Δ; sst2Δ* | pDAM194 | SB9 | This study |
| SB37 | *MAT**a**/**a**; ura3Δ; his3Δ; sst2Δ; ste3Δ; pPGK1-GPA1-tCYC1-X-3; gpa1Δ0* | pDAM194 | SB30 | This study |
| SB38 | *MAT**a**/**a**; ura3Δ; his3Δ; sst2Δ; ste3Δ; pRNR2-GPA1-tCYC1-X-3; gpa1Δ0* | pDAM194 | SB24 | This study |
| SB39 | *MAT**a**/**a**; ura3Δ; his3Δ; pPGK1-GPA1-tCYC1-X-3; gpa1Δ0* | pDAM194 | SB35 | This study |

| | | | | |
|---|---|---|---|---|
| SB40 | *MAT**a**/**a**; ura3Δ; his3Δ; pRNR2-GPA1-tCYC1-X-3; gpa1Δ0* | pDAM194 | SB26 | This study |
| SB41 | *MAT**a**/**a**; ura3Δ; his3Δ; sst2Δ; ste3Δ; pPGK1-GPA1-tCYC1-X-3; gpa1Δ0; ste2Δ* | pDAM194 | SB31 | This study |
| SB42 | *MAT**a**/**a**; ura3Δ; his3Δ; pPGK1-GPA1-tCYC1-X-3; gpa1Δ0; ste2Δ* | pDAM194 | SB33 | This study |
| SB45 | *MAT**a**/**a**; ura3Δ; his3Δ; sst2Δ; ste3Δ; pPGK1-GPA1-tCYC1-X-3; gpa1Δ0; ste2Δ* | pDAM194+pDAM71 | SB41 | This study |
| SB46 | *MAT**a**/**a**; ura3Δ; his3Δ; sst2Δ; ste3Δ; pPGK1-GPA1-tCYC1-X-3; gpa1Δ0; ste2Δ* | pDAM194+pDAM72 | SB41 | This study |
| SB47 | *MAT**a**/**a**; ura3Δ; his3Δ; sst2Δ; ste3Δ; pPGK1-GPA1-tCYC1-X-3; gpa1Δ0; ste2Δ* | pDAM194+pDAM74 | SB41 | This study |
| SB48 | *MAT**a**/**a**; ura3Δ; his3Δ; pPGK1-GPA1-tCYC1-X-3; gpa1Δ0; ste2Δ* | pDAM194+pDAM71 | SB42 | This study |
| SB49 | *MAT**a**/**a**; ura3Δ; his3Δ; pPGK1-GPA1-tCYC1-X-3; gpa1Δ0; ste2Δ* | pDAM194+pDAM72 | SB42 | This study |
| SB50 | *MAT**a**/**a**; ura3Δ; his3Δ; pPGK1-GPA1-tCYC1-X-3; gpa1Δ0; ste2Δ* | pDAM194+pDAM74 | SB42 | This study |

### Supplementary Table S5

| Plasmid | Design | Plasmid marker | Backbone | Origin of replication | Reference |
|---|---|---|---|---|---|
| pEDJ391 | pTEF1-*Cas9*-tCYC1 | *LEU2* | N/A | CEN/ARS | Jensen et al. 2021 |
| pEDJ400 | USER cloning site | *URA3* | N/A | 2µ | Jensen et al. 2021 |
| pEDJ437 | USER cloning site | *HIS3* | N/A | 2µ | Jensen et al. 2021 |
| pDAM1 | gRNA_*STE2* (TACAGTTTGAAACCA AACCA) | *NatR* | pCfB3050 | 2µ | This study |
| pDAM2 | gRNA_*STE3* (GATATTGGTGTACCT ACACA) | *NatR* | pCfB3050 | 2µ | This study |
| pDAM3 | gRNA_*FUS1* (AGAAGTATAACGACA CCCAG) | *NatR* | pCfB3050 | 2µ | This study |
| pDAM4 | gRNA_*SST2* (ATCCCAACCACGCTTG GACA) | *NatR* | pCfB3050 | 2µ | This study |
| pDAM6 | gRNA_*GPA1-C-term* (GTTTTGCTGGATGATT AGAT) | *NatR* | pCfB3050 | 2µ | This study |
| pDAM7 | gRNA_*STE2* (TACAGTTTGAAACCA AACCA) | *URA3* | pEDJ400 | 2µ | This study |

| | | | | | |
|---|---|---|---|---|---|
| pDAM8 | gRNA_*STE3* (GATATTGGTGTACCTACACA) | *HIS3* | pEDJ437 | 2µ | This study |
| pDAM22 | gRNA_X-4_XI-3_XII-4 | *NatR* | pTAJAK-71 | 2µ | This study |
| pDAM23 | *tADH1-RnPTS<-pTEF1-pPGK1->RnSPR-tCYC1* (X-4) | Markerfree | pCfB3035 | N/A | This study |
| pDAM24 | *tADH1-PaPCBD1<-pTEF1-pPGK1->RnDHPR-tCYC1* (XII-4) | Markerfree | pCfB3040 | N/A | This study |
| pDAM25 | *tADH1-HsASMT<-pTEF1-pPGK1->BtAANAT-tCYC1* (X-2) | Markerfree | pCfB2899 | N/A | This study |
| pDAM30 | pPGK1-*GPA1*-tCYC1 | *LEU2* | pRS415U | CEN/ARS | This study |
| pDAM32 | pPGK1-*GPA1/Gαi2*-tCYC1 | *LEU2* | pRS415U | CEN/ARS | This study |
| pDAM47 | pRNR2-*GPA1*-tCYC1 | *LEU2* | pRS415U | CEN/ARS | This study |
| pDAM49 | pRNR2-*GPA1/Gαi2*-tCYC1 | *LEU2* | pRS415U | CEN/ARS | This study |
| pDAM50 | pALD6-*GPA1*-tCYC1 | *LEU2* | pRS415U | CEN/ARS | This study |
| pDAM52 | pALD6-*GPA1/Gαi2*-tCYC1 | *LEU2* | pRS415U | CEN/ARS | This study |

| | | | | | |
|---|---|---|---|---|---|
| pDAM71 | pCCW12:*ADORA2B*:tCYC1 | *HIS3* | pEDJ437 | 2μ | This study |
| pDAM72 | pCCW12:*MTNR1A*:tCYC1 | *HIS3* | pEDJ437 | 2μ | This study |
| pDAM74 | pCCW12:*MAM2*:tCYC1 | *HIS3* | pEDJ437 | 2μ | This study |
| pDAM76 | pCCW12:*HT4R*:tCYC1 | *HIS3* | pEDJ437 | 2μ | This study |
| pDAM77 | gRNA_*GPA1* (GCATCACATCAATAATCCAG) - promoter target | *NatR* | pDAM1 | 2μ | This study |
| pDAM78 | gRNA_*GPA1* (CTGTTTCCGAAGATGCAAGA) - terminator target | *NatR* | pDAM1 | 2μ | This study |
| pDAM82 | gRNA_*GPA1*_promoter+terminator target | *HIS3* | pEDJ437 | 2μ | This study |
| pDAM122 | pTEF1-*mKate2*-tCYC1 | Markerfree | pCfB3039 | N/A | This study |
| pDAM123 | pTEF1-*α-leader_P-factor*-tCYC1 | Markerfree | pCfB2899 | N/A | This study |
| pDAM182 | gRNA_*MATα* (CAAATCATACAGAAACACAG) | *HIS3* | pEDJ437 | 2μ | This study |

| | | | | | |
|---|---|---|---|---|---|
| pDAM194 | pFUS1-*yEGFP*-tFUS1 | *URA3* | pJV452 | CEN/ARS | This study |
| pDAM215 | pTEF1-*Cas9*-tCYC1 | *URA3* | pRS416U | CEN/ARS | This study |
| pDAM216 | pCCW12:*ADRB2*: tCYC1 | *HIS3* | pEDJ437 | 2µ | This study |
| pDAM217 | pCCW12*:CXCR4:tCYC1* | *HIS3* | pEDJ437 | 2µ | This study |
| pDAM218 | pCCW12:*GLP-1R*: tCYC1 | *HIS3* | pEDJ437 | 2µ | This study |
| pDAM219 | pCCW12:*CaSTE2*: tCYC1 | *HIS3* | pEDJ437 | 2µ | This study |
| pDAM220 | pCCW12:*FgSTE2*: tCYC1 | *HIS3* | pEDJ437 | 2µ | This study |
| pDAM221 | pCCW12:*ZtSTE2:* tCYC1 | *HIS3* | pEDJ437 | 2µ | This study |
| pDAM222 | pCCW12:*TrSTE2:* tCYC1 | *HIS3* | pEDJ437 | 2µ | This study |
| pDAM223 | pCCW12:*MsSTE3:* tCYC1 | *HIS3* | pEDJ437 | 2µ | This study |
| ID6911 | gRNA_X-3 | *URA3* | N/A | 2µ | This study |
| PL_12_I3 | gRNA_*MATa* (ATCAATATCACCCCAAGCAC) | *URA3* | N/A | 2µ | This study |

| | | | | | |
|---|---|---|---|---|---|
| pJV452 | USER cloning site | *URA3* | pEDJ400 | CEN/ARS | This study |
| pCfB2899 | USER integration site (X-2) | Markerfree | N/A | N/A | Jessop-Fabre et al. 2016 |
| pCfB3020 | gRNA_X-2 | *NatR* | N/A | 2µ | Jessop-Fabre et al. 2016 |
| pCfB3035 | USER integration site (X-4) | Markerfree | N/A | N/A | Jessop-Fabre et al. 2016 |
| pCfB3039 | USER integration site (XII-2) | Markerfree | N/A | N/A | Jessop-Fabre et al. 2016 |
| pCfB3040 | USER integration site (XII-4) | Markerfree | N/A | N/A | Jessop-Fabre et al. 2016 |
| pCfB3042 | gRNA_X-4 | *NatR* | N/A | 2µ | Jessop-Fabre et al. 2016 |
| pCfB3044 | gRNA_XI-2 | *NatR* | N/A | 2µ | Jessop-Fabre et al. 2016 |
| pCfB3045 | gRNA_XI-3 | *NatR* | N/A | 2µ | Jessop-Fabre et al. 2016 |
| pCfB3048 | gRNA_XII-2 | *NatR* | N/A | 2µ | Jessop-Fabre et al. 2016 |
| pCfB3049 | gRNA_XII-4 | *NatR* | N/A | 2µ | Jessop-Fabre et al. 2016 |

| | | | | | |
|---|---|---|---|---|---|
| pCfB3050 | gRNA_XII-5 | *NatR* | N/A | 2µ | Jessop-Fabre et al. 2016 |
| pCfB9221 | *tADH1-HsDDC<-pTDH3-pTEF1->SmTPH-tCYC1* (XI-3) | Markerfree | N/A | N/A | Gift from Dr. Nicholas Milne |
| pCfB1248 | *pX-4-LoxP-SpHIS5-PaPCBD1<-pTEF1-pPGK1->RnDHPR* | *SpHIS5* | N/A | N/A | Germann et al. 2016 |
| pCfB1251 | *pX-3-LoxP-KlLEU2-RnPTS<-pTEF1-pPGK1->RnSPR* | *KILEU2* | N/A | N/A | Germann et al. 2016 |
| pCfB1252 | *pXII-1-LoxP-KILEU2-HsASMT<-pTEF1* | *KILEU2* | N/A | N/A | Germann et al. 2016 |
| pCfB2772 | *Ty2::KlURA3-pPGK1-SmTPH-tCYC1* | N/A | N/A | N/A | Germann et al. 2016 |
| pCfB2628 | *pXI-5-LoxP-SpHIS5-HsDDC<-pTEF1-pPGK1->BtAANAT* | *SpHIS5* | N/A | N/A | Germann et al. 2016 |
| pTAJAK-71 | USER cloning site | *NatR* | N/A | 2µ | Jakočiūnas et al. 2015 |
| pRS416U | USER cloning site | *URA3* | N/A | CEN/ARS | Jensen et al. 2017 |
| pRS415U | USER cloning site | *LEU2* | N/A | CEN/ARS | Jensen et al. 2017 |

| | | | | | |
|---|---|---|---|---|---|
| pYR11 | pTDH3:his6-*mKate2*-*MBP*-tIDP1 (XII-5) | N/A | N/A | N/A | Romero-Suarez et al. 2021 |
| pDD110 | pCCW12:*Cas9*:tENO2: pSNR52: *GFP*_dropout:tSUP1 | *NatR* | N/A | CEN/ARS | This study |
| pDD111 | gRNA_*URA3* (TAGCGGTTTGAAGCA GGCGG) | *NatR* | DD110 | CEN/ARS | This study |
| pDD115 | gRNA_*HIS3* (CATTTGTAATACGCTT TACT) | *NatR* | DD110 | CEN/ARS | This study |
| pDD118 | pTEF1:*Cas9*:tENO2 | *NatR* | N/A | CEN/ARS | Durmusoglu et al. 2021 |

### Supplementary Table S6

| Oligo Name | Oligo Sequence |
|---|---|
| DAM1 | TACAGTTTGAAACCAAACCAGTTTTAGAGCTAGAAATAGCAAG |
| DAM2 | GATATTGGTGTACCTACACAGTTTTAGAGCTAGAAATAGCAAG |
| DAM3 | AGAAGTATAACGACACCCAGGTTTTAGAGCTAGAAATAGCAAG |
| DAM4 | ATCCCAACCACGCTTGGACAGTTTTAGAGCTAGAAATAGCAAG |
| DAM7 | GTGGCTACTCTGATTAGTATG |
| DAM8 | CGAAGGTCACGAAATTACTTTTTCAAAGCCGTAAATTTTGATTTTGATTCTTGGATATGG |
| DAM9 | ACTTAAAAATGCACCGTTAAGAACCATATCCAAGAATCAAAATCAAAATTTACGGCTTTG |
| DAM10 | GTTATCTCATTAGTAATCC |
| DAM13 | GATTCGCCAAAGTATTCTTACC |
| DAM14 | TACTCCTAGTCCAGTAAATATAATGCGACACTCTTGTGGAAAATTTTGATAGTATTTTGC |
| DAM15 | TGTAGGAAAGGCAAAATACTATCAAAATTTTCCACAAGAGTGTCGCATTATATTTACTGG |
| DAM16 | GAATATACTGACGTATCCATTG |
| DAM19 | GCAAGTTATTACAGTCTTAC |
| DAM20 | AATTGTACCTGAAGATGAGTAAGACTCTCAATGAAACTTACAACTCTATCTTTAATTACC |
| DAM21 | GTTATAGGTTCAATTTGGTAATTAAAGATAGAGTTGTAAGTTTCATTGAGAGTCTTACTC |
| DAM22 | GCAAGAAAACACACTACCTG |
| DAM31 | CGTGTCGGCGATGCAGAAGCG |
| DAM34 | CCCGAGTATTTACCAGGGAG |
| DAM37 | GAGTAGTTCGTACTGTTTAAGGTTTTGCTGGATGATCAAATCCGTGACTGCACTCAATAC |
| DAM38 | TGATCATCCAGCAAAACCTTAAACAGTACGAACTACTCTGAATCGCGTGCATTCATCCGC |
| DAM39 | CATCTACACAATTAGCAAGG |
| DAM40 | TAAAAAAGGAGTAGAAACATTTTGAAGCTATGGTGTGTGCcataacgcgttacacggaag |
| DAM41 | ccctccttactgctctccttccgtgtaacgcgttatgGCACACACCATAGCTTCAAAATG |
| DAM42 | GATTAACTCCTCTCCTTTGGACACCATTGTTTTTTGTAATTAAAACTTAGATTAGATTGC |
| DAM43 | AGCATAGCAATCTAATCTAAGTTTTAATTACAAAAAACAATGGTGTCCAAAGGAGAGGAG |
| DAM44 | aattcgcttatttagaagtgtcaacaacgtatctacTTACTTATACAATTCATCCATACC |
| DAM45 | CTGGCTTAGGCGGTGGTATGGATGAATTGTATAAGTAAgtagatacgttgttgacacttc |
| DAM46 | gcagtacaaggacgcgttaagaaaaatttcgagagagtcggagcgacctcatgctatacc |
| DAM47 | ggtatagcatgaggtcgctccgactctctcgaaatttttcttaacgcgtccttgtactgc |
| DAM48 | AGCGAACGTAAGAGAGG |

| | |
|---|---|
| DAM53 | CATCAACAACAGGGTCAGCAG |
| DAM54 | ACAACACCAGTGAATAATTCTTCACCTTTAGACATTGTTTTTTTGATTTTCAGAAACTTG |
| DAM55 | TGCTGGTATTACCCATGGTATGGATGAATTGTACAAATAATGAAAATAATATTGACGTTC |
| DAM56 | CTGAAGTATCCTATATCAAC |
| DAM58 | CGTGCGAUCGAAGAAGTACCTTCAAAGAATGG |
| DAM59 | ATGACAGAUATCCGCTCTAACCGAAAAGG |
| DAM63 | ACCTGCACUAAAACAATGTTGTTGGAAACACAAGATGCG |
| DAM64 | ATCTGTCAUTCAGGATCCGAGTCCAACGCC |
| DAM65 | ACCTGCACUAAAACAATGCAAGGTAATGGTTC |
| DAM66 | ATCTGTCAUTCAGGATCCAACGGAGTC |
| DAM67 | ACCTGCACUAAAACAATGAGGCAACCTTGGTGG |
| DAM68 | ATCTGTCAUTCAGGATCCCGTCCAC |
| DAM76 | ATCTGTCAUAAAACAATGAGATTTCCTTC |
| DAM77 | ACCTGCACUAAAACAATGAGGCAGGTTTGG |
| DAM78 | ACCTGCACUTTACAAATTGGGCCTATCTGG |
| DAM80 | CACGCGAUAGTGCAGGUATCCGCTCTAACCGAAAAGG |
| DAM81 | TTCTCAAGCAAGGTTTTCAG |
| DAM85 | GTCTCTATTGGAAAACTGAATGC |
| DAM115 | CACGCGAUTTACAAATTGGGCCTATCTGG |
| DAM116 | AGTGCAGGUGGTAATTGGACAAATAAATACGTGTATTAAG |
| DAM117 | CGTGCGAUGAAACTAAACTAAAAACCGTAC |
| DAM118 | AGTGCAGGUTGTATTCTGATAGTATG |
| DAM160 | ACCTGCACUAAAACAATGGATAAATTGGATGCTAATGTTTC |
| DAM162 | ATCTGTCAUCCTCTCAATAGGGAAAACTGGCC |
| DAM191 | CGTGCGAUGCGGCCGCAACCAGGGCAAAGCAAAATAAAAG |
| DAM192 | AGTGCAGGUTATTGATATAGTGTTTAAGCG |
| DAM209 | caataggcaagaagtaggcgagagccgacatacgagactAGCCACAGCTGTGCATTACGC |
| DAM210 | aataggcaagaagtaggcgagagccgacatacgagactGAAACTAAACTAAAAACCGTAC |
| DAM211 | acaataggcaagaagtaggcgagagccgacatacgagactCGAAGAAGTACCTTCAAAG |
| DAM212 | cttttactagcatatcaatatccgtttcattgaaaagtggTTCTCAAGCAAGGTTTTCAG |
| DAM218 | GCATCACATCAATAATCCAGGTTTTAGAGCTAGAAATAGCAAG |
| DAM219 | CTGTTTCCGAAGATGCAAGAGTTTTAGAGCTAGAAATAGCAAG |

| | |
|---|---|
| DAM235 | GGAGAAGAAAAATAGTAATTTTTTCTGGTGCTGCTGCTCCTTCTGTGATGCTAAAATACC |
| DAM236 | AGTAGAAGCTATTCATACTGTAAATTGGTATTTTAGCATCACAGAAGGAGCAGCAGCACC |
| DAM288 | ATCTGTCAUAAAACAATGGTTTCTGAACTCATCAAGG |
| DAM289 | CACGCGAUTTATCTGTGTCCCAACTTAGATGG |
| DAM335 | ACCTGCACUAAAACAGAATTCAACGTTGGATC |
| DAM336 | ATCTGTCAUTTATAGAAGAGAGTCGTTGG |
| DAM337 | ACCTGCACUAAAACAATGTCCATACCGTTGCCGCTTC |
| DAM338 | ATCTGTCAUTCAGGAGCTGTGAAAAGAGG |
| DAM339 | ACCTGCACUAAAACAATGGCCGGAGCACCCG |
| DAM340 | ATCTGTCAUTCATGAGCAAGAGGCCTGAC |
| DAM341 | ACCTGCACUAAAACAATGAACATTAATTCTACGTTCATACC |
| DAM342 | ATCTGTCAUCTATACACGTTTGATTGTAATCTGC |
| DAM343 | ACCTGCACUAAAACAATGTCCAAAGAAGTATTCGACC |
| DAM344 | ATCTGTCAUCTACAAGGGGGCACGTATCCTCTCCTCTCTCTG |
| DAM345 | ACCTGCACUAAAACAATGGTAGTTACCGCACCTCCC |
| DAM346 | ATCTGTCAUCTAATCGGACCTTACTGAGTAAGAACGCCC |
| DAM347 | ACCTGCACUAAAACAATGGCCCCACACTTTG |
| DAM348 | ATCTGTCAUCAATCTGTTACTAGTGACGCTGAAATTACG |
| DAM349 | ACCTGCACUAAAACAATGTCAGGCTTTGCTAC |
| DAM350 | ATCTGTCAUAGCGAAACGATTAGTTTTCGTGTCATC |
| DAM418 | TGATCATCCAGCAAAACCTTAAACTATGTGGCCTTATCTGAATCGCGTGCATTCATCCGC |
| DAM419 | GATAAGGCCACATAGTTTAAGGTTTTGCTGGATGATCAAATCCGTGACTGCACTCAATAC |
| DAM422 | TGATCATCCAGCAAAACCTTAAAGATTCAGGGATTTTGTGAATCGCGTGCATTCATCCGC |
| DAM423 | CAAAATCCCTGAATCTTTAAGGTTTTGCTGGATGATCAAATCCGTGACTGCACTCAATAC |
| DAM426 | TGATCATCCAGCAAAACCTTAAAGAAACGGGGTTCTTGTGAATCGCGTGCATTCATCCGC |
| DAM427 | CAAGAACCCCGTTTCTTTAAGGTTTTGCTGGATGATCAAATCCGTGACTGCACTCAATAC |
| DAM485 | CAAATCATACAGAAACACAGGTTTTAGAGCTAGAAATAGCAAG |
| DAM550 | TGATCATCCAGCAAAACCTTAAAATGTGTGGGTTGATTTGAATCGCGTGCATTCATCCGC |
| DAM551 | AATCAACCCACACATTTTAAGGTTTTGCTGGATGATCAAATCCGTGACTGCACTCAATAC |
| DAM594 | 5/PHOS/GATCATTTATCTTTCACTGCGGAGAAG |
| EDJ134 | CGTGCGAUAGCCACAGCTGTGCATTACG |
| EDJ318 | AGTGCAGGUTGTTTTATATTTGTTGTAAAAAGTAG |

| | |
|---|---|
| EDJ325 | CACGCGAUTCACACCTTCCTCTTCTTCTTGG |
| 1564 | CGTGCGAUGCACACACCATAGCTTC |
| 1565 | ATGACAGAUTTGTAATTAAAACTTAG |
| TJOS-62 | CGTGCGAUagggaacaaaagctggagct |
| TJOS-63 | AGTGCAGGUagggaacaaaagctggagct |
| TJOS-64 | ATCTGTCAUagggaacaaaagctggagct |
| TJOS-65 | CACGCGAUtaactaattacatgactcga |
| TJOS-66 | ACCTGCACUtaactaattacatgactcga |
| TJOS-67 | ATGACAGAUtaactaattacatgactcga |
| MAB1 | CGTGCGAUGGCAACATAGCAGTACGTCGCCAAA |
| ID904 | CCGTGCAATACCAAAATCG |
| JV498 | ttacaggcaagcgatccgtcACCTGCACTCAACGGAATGC |
| JV499 | tttatagcacgtgatactccTTCAGGTGGCACTTTTCGGG |
| JV500 | CCCGAAAAGTGCCACCTGAAggagtatcacgtgctataaaaataatt |
| JV501 | GCATTCCGTTGAGTGCAGGTgacggatcgcttgcctgtaa |
| JV502 | CGTGCGAUAATAATCAGAACTCCAACAATAGTCA |
| JV503 | CACGCGAUATTCACCAGACCCGCTCCT |
| JZ1 | ctaatgtgtccgcgtttctaGTTTTAGAGCTAGAAATAGCAAG |
| MAD1 | ATCAATATCACCCCAAGCACGTTTTAGAGCTAGAAATAGCAAG |
| PR_26_D7 | TGTTACACTCTCTGGTAACTTAG |
| PR_26_D8 | AAGATAAGAACAAAGAATGATGCTAAG |
| PR_26_D8 | AAGATAAGAACAAAGAATGATGCTAAG |
| DDpr097 | GCCGCCGTTGTTGTTTTT |
| DDpr098 | TCATAACACAGTCCTTTCCCGC |
| DDpr254 | GGTTCTGGCGAGGTATTGGATAGTTC |
| DDpr255 | TGGCTGTGGTTTCAGGGTCCATAAAGC |
| DDpr262 | gatcTAGCGGTTTGAAGCAGGCGG |
| DDpr263 | aaacCCGCCTGCTTCAAACCGCTA |
| DDpr264 | gatcCATTTGTAATACGCTTTACT |
| DDpr265 | aaacAGTAAAGCGTATTACAAATG |
| DDpr273 | CGACGTTGAAATTGAGGCTAC |

| | |
|---|---|
| DDpr274 | CTACACGTTCGCTATGCTTC |
| DDpr276 | GCCGCCGTTGTTGTTATTGT |
| DDpr277 | CTGTTCCCTAGCATGTACGT |
| 5 | acctgcacuttgtaattaaaacttag |
| 8 | acctgcacuttgtaattaaaacttag |
| 350 | CGTGCGAUTTATTCTCCTTTGTAGACCACAAT |
| 389 | CACGCGAUTTAAATGTCATAGAAGTCCACGTG |
| 1564 | CGTGCGAUGCACACACCATAGCTTC |
| 1565 | CGTGCGAUGCACACACCATAGCTTC |
| 1761 | atctgtcaUaaaacaatgagcaccccgagcattcattg |
| 1762 | CACGCGAUTTAACGATCGCTATTACGACGCAGTG |
| 2254 | AGTGCAGGUAAAACAATGGGTAGCAGCGAAGATC |
| 2255 | CGTGCGAUTTATTTACGTGCCAGGATTGCATC |
| 2149 | CGTGCGAUTTACTTTCTACCTTCAGCAG |
| 2153 | CACGCGAUTTAGAAGTAAGCTGGAGTC |

### Supplementary Table S7

| ID | Insert | Sequence |
|---|---|---|
| gDAM1 | *GPA1/Gαi2* | AAAACAATGGGGTGTACAGTGAGCACGCAAACAATAGGAGACGAAAG<br>TGATCCTTTTCTACAGAACAAAAGAGCCAATGATGTCATCGAGCAATC<br>GTTGCAGCTGGAGAAACAACGTGACAAGAATGAAATAAAACTGTTACT<br>ATTAGGTGCCGGTGAGTCAGGTAAATCAACGGTTTTAAAACAATTAAA<br>ATTATTACATCAAGGCGGTTTCTCCCATCAAGAAAGGTTACAGTATGC<br>TCAAGTGATACGGGCAGATGCCATACAATCAATGAAAATTTTGATTATT<br>CAGGCCAGAAAACTAGGTATTCAACTTGACTGTGATGATCCGATCAAC<br>AATAAAGATTTGTTTGCATGTAAGAGAATACTGCTAAAGGCTAAAGCTT<br>TAGATTATATCAACGCCAGTGTTGCCGGTGGTTCTGATTTTCTAAATG<br>ATTATGTACTGAAGTACTCAGAAAGGTATGAAACTAGGAGGCGTGTTC<br>AGAGTACCGGACGAGCAAAAGCTGCTTTCGATGAAGACGGAAATATT<br>TCTAATGTCAAAAGTGACACTGACAGAGATGCTGAAACGGTGACGCA<br>AAATGAGGATGTTGATAGAAACAACAGTAGTAGAATTAACCTACAGGA<br>TATTTGCAAGGACTTGAACCAAGAAGGCGATGACCAGATGTTTGTTAG<br>AAAAACATCAAGGGAAATTCAAGGACAAAATAGACGAAATCTTATTCA<br>CGAAGACATTGCTAAGGCAATAAAGCAACTTTGGAATAACGACAAAGG<br>TATAAAGCAGTGCTTTGCACGTTCTAATGAGTTTCAATTGGAGGGCTC<br>AGCTGCATACTACTTTGATAACATTGAGAAATTTGCTAGTCCGAATTAT<br>GTCTGTACGGATGAAGACATTTTGAAGGGCCGTATAAAGACTACAGG<br>CATTACAGAAACCGAATTTAACATCGGCTCGTCCAAATTCAAGGTTCT<br>CGACGCTGGTGGGCAGCGTTCTGAACGTAAGAAGTGGATTCATTGTT<br>TCGAAGGAATTACAGCAGTTTTATTTGTTTTAGCAATGAGTGAATACGA<br>CCAGATGTTGTTTGAGGATGAAAGAGTGAACAGAATGCATGAATCAAT<br>AATGCTATTTGACACGTTATTGAACTCTAAGTGGTTCAAAGATACACC<br>GTTTATTTTGTTTTTAAATAAAATTGATTTGTTCGAGGAAAAGGTAAAAA<br>GCATGCCCATAAGAAAGTACTTTCCTGATTACCAGGGACGTGTCGGC<br>GATGCAGAAGCGGGTCTAAAATATTTTGAGAAGATATTTTTGAGCTTG<br>AATAAGACAAACAAACCAATCTACGTGAAACGAACCTGCGCTACCGAT<br>ACCCAAACTATGAAGTTCGTATTGAGTGCAGTCACGGATTTGATCATC<br>CAGCAAAACCTTAAAGATTGCGGCCTGTTCTGA |

| | | |
|---|---|---|
| gDAM2 | *ADORA2B* | TTGTTGGAAACACAAGATGCGTTGTATGTTGCGTTGGAATTGGTAATT<br>GCTGCGCTATCGGTTGCGGGAAATGTTTTGGTTTGTGCTGCGGTTGG<br>AACGGCGAATACCTTGCAAACGCCTACTAATTATTTTTTGGTTTCATTG<br>GCAGCGGCTGATGTTGCTGTTGGACTATTTGCTATTCCTTTCGCTATT<br>ACTATTTCTTTGGGATTTTGTACCGATTTTTATGGATGTCTATTTCTAG<br>CTTGTTTTGTTTTGGTTCTAACGCAATCTTCAATTTTTTCTCTATTGGCT<br>GTTGCCGTAGATAGGTATTTGGCTATTTGCGTACCGCTAAGGTACAAG<br>TCCCTTGTAACGGGAACTCGTGCCAGGGGAGTAATAGCAGTACTATG<br>GGTACTAGCTTTCGGAATTGGACTTACCCCTTTTTTGGGATGGAATTC<br>CAAGGATTCCGCTACTAATAATTGTACAGAGCCTTGGGACGGAACTAC<br>GAACGAGTCTTGTTGTCTAGTTAAATGCCTATTCGAAAACGTTGTACC<br>TATGTCTTATATGGTGTACTTTAACTTTTTCGGATGCGTGTTGCCTCCT<br>TTGCTAATCATGTTGGTTATTTATATAAAAATTTTTTTGGTTGCTTGTAG<br>GCAACTACAACGTACCGAATTGATGGATCATTCGAGGACTACTCTACA<br>AAGGGAAATTCACGCCGCTAAATCCTTGGCTATGATAGTTGGAATATT<br>CGCTTTGTGTTGGCTCCCTGTTCACGCAGTGAATTGCGTAACCCTATT<br>TCAACCTGCACAAGGCAAGAACAAACCTAAATGGGCCATGAACATGG<br>CTATACTATTGTCCCACGCTAACTCCGTGGTAAACCCTATAGTATACG<br>CATATAGGAATCGTGATTTTCGTTATACCTTCCATAAGATAATTTCAAG<br>GTACCTACTATGTCAGGCCGACGTAAAATCCGGAAACGGCCAAGCAG<br>GAGTGCAACCTGCACTAGGCGTTGGACTCGGATCC |

| | | |
|---|---|---|
| gDAM3 | *MTNR1A* | AAAACAATGCAAGGTAATGGTTCTGCTTTGCCAAATGCTTCTCAACCA<br>GTTTTGAGAGGTGATGGTGCTAGACCTTCTTGGTTGGCTTCTGCTTTA<br>GCTTGTGTTTTGATTTTCACCATCGTTGTCGATATCTTGGGTAACTTGT<br>TGGTTATCTTGTCCGTCTACCGTAACAAGAAATTGAGAAACGCTGGTA<br>ACATCTTCGTTGTTTCTTTGGCTGTTGCTGATTTGGTTGTTGCTATCTA<br>TCCATATCCACTGGTCTTGATGTCCATTTTTAACAACGGTTGGAACTTG<br>GGTTACTTGCATTGTCAAGTTTCTGGTTTCTTGATGGGTTTGTCCGTTA<br>TTGGTTCCATTTTCAACATTACCGGTATCGCCATTAACAGGTACTGTTA<br>CATTTGCCACTCACTGAAGTACGACAAGTTGTACTCTTCTAAGAACTC<br>CTTGTGCTACGTTTTGTTGATCTGGTTGTTAACTTTGGCTGCTGTTTTG<br>CCTAATTTGAGAGCTGGTACATTGCAATACGATCCAAGAATCTACTCT<br>TGTACCTTCGCTCAATCTGTTTCTTCTGCTTACACTATTGCCGTTGTCG<br>TTTTCCATTTTTTGGTCCCAATGATTATCGTCATCTTCTGCTACTTGAG<br>AATCTGGATTTTGGTCTTGCAAGTCAGACAAAGAGTTAAGCCAGATAG<br>AAAGCCAAAATTGAAGCCACAAGACTTCAGAAACTTCGTTACCATGTT<br>TGTGGTTTTCGTTTTGTTCGCTATTTGTTGGGCTCCATTGAACTTTATT<br>GGTTTGGCAGTTGCTTCTGATCCAGCTTCTATGGTTCCAAGAATTCCA<br>GAATGGTTGTTCGTTGCTTCTTACTACATGGCTTACTTCAACTCTTGTT<br>TGAACGCCATTATCTACGGCTTGTTGAATCAGAACTTTAGGAAAGAGT<br>ACAGGCGTATCATCGTTTCTTTGTGTACTGCTAGAGTTTTCTTCGTCG<br>ATTCCTCTAATGATGTTGCCGATAGAGTTAAGTGGAAACCATCTCCAT<br>TGATGACCAACAACAATGTTGTCAAGGTTGACTCCGTTGGATCCTGA |

| | | |
|---|---|---|
| gDAM4 | *MAM2* | AAAACAATGAGGCAACCTTGGTGGAAAGACTTTACTATTCCCGATGCA<br>AGCGCAATTATTCACCAAAATATTACCATTGTGTCTATTGTCGGAGAG<br>ATTGAAGTGCCTGTTTCAACAATTGATGCATATGAAAGGGATAGGTTA<br>TTAACTGGAATGACTTTATCTGCCCAATTAGCTTTAGGAGTGTTAACCA<br>TTTTAATGGTTTGTCTGTTATCATCAAGCGAAAAACGTAAACACCCTGT<br>TTTTGTTTTTAATTCGGCAAGTATTGTTGCAATGTGTTTACGCGCCATT<br>TTAAATATCGTGACCATCTGCTCGAATTCGTACAGTATCCTCGTTAATT<br>ACGGGTTTATCTTAAACATGGTTCATATGTATGTGCATGTGTTTAATAT<br>TTTAATTTTATTATTAGCACCGGTGATCATTTTTACTGCTGAGATGTCG<br>ATGATGATTCAAGTTAGGATCATTTGTGCACATGATAGGAAAACACAA<br>CGTATCATGACTGTTATTAGTGCCTGCTTAACTGTTTTAGTTCTCGCAT<br>TTTGGATTACTAACATGTGTCAACAGATTCAGTATCTCTTATGGTTAAC<br>TCCTTTATCGTCGAAAACCATTGTTGGATACTCTTGGCCCTACTTTATT<br>GCTAAAATCTTATTTGCTTTTTCGATTATTTTTCACAGTGGTGTTTTTTC<br>ATACAAACTCTTTAGGGCCATCTTAATCCGCAAAAAAATTGGGCAATTT<br>CCTTTTGGTCCGATGCAGTGTATTTTAGTTATTTCGTGCCAATGTTTAA<br>TTGTTCCTGCTACCTTTACTATCATCGATAGTTTTATCCATACGTATGA<br>TGGCTTTTCGTCTATGACTCAATGTCTCCTGATCATTTCTTTACCATTA<br>TCGAGTTTATGGGCGTCTAGTACAGCTCTCAAATTACAATCGATGAAA<br>ACTTCATCTGCGCAAGGAGAAACCACCGAGGTTTCGATTAGGGTTGA<br>TAGGACGTTTGATATCAAACATACTCCCAGTGACGATTATTCGATTTCT<br>GATGAATCTGAAACTAAAAAGTGGACGGGATCCTGA |

| | | |
|---|---|---|
| gDAM5 | *HT4R* | AAAACAATGGATAAATTGGATGCTAATGTTTCTTCTGAAGAAGGTTTTG<br>GTTCTGTTGAAAAAGTTGTTTTGTTGACTTTTTTGTCTACTGTTATTTTG<br>ATGGCTATTTTGGGTAATTTGTTGGTTATGGTTGCTGTTTGTTGGGATA<br>GACAATTGAGAAAAATTAAAACTAATTATTTTATTGTTTCTTTGGCTTTT<br>GCTGATTTGTTGGTTTCTGTTTTGGTTATGCCATTTGGTGCTATTGAAT<br>TGGTTCAAGATATTTGGATTTATGGTGAAGTTTTTTGTTTGGTTAGAAC<br>TTCTTTGGATGTTTTGTTGACTACTGCTTCTATTTTTCATTTGTGTTGTA<br>TTTCTTTGGATAGATATTATGCTATTTGTTGTCAACCATTGGTTTATAGA<br>AATAAAATGACTCCATTGAGAATTGCTTTGATGTTGGGTGGTTGTTGG<br>GTTATTCCAACTTTTATTTCTTTTTTGCCAATTATGCAAGGTTGGAATAA<br>TATTGGTATTATTGATTTGATTGAAAAAAGAAAATTTAATCAAAATTCTA<br>ATTCTACTTATTGTGTTTTTATGGTTAATAAACCATATGCTATTACTTGT<br>TCTGTTGTTGCTTTTTATATTCCATTTTTGTTGATGGTTTTGGCTTATTA<br>TAGAATTTATGTTACTGCTAAAGAACATGCTCATCAAATTCAAATGTTG<br>CAAAGAGCTGGTGCTTCTTCTGAATCTAGACCACAATCTGCTGATCAA<br>CATTCTACTCATAGAATGAGAACTGAAACTAAAGCTGCTAAAACTTTGT<br>GTATTATTATGGGTTGTTTTTGTTTGTGTTGGGCTCCATTTTTTGTTACT<br>AATATTGTTGATCCATTTATTGATTATACTGTTCCAGGTCAAGTTTGGA<br>CTGCTTTTTTGTGGTTGGGTTATATTAATTCTGGTTTGAATCCATTTTT<br>GTATGCTTTTTTGAATAAATCTTTTAGAAGAGCTTTTTTGATTATTTTGT<br>GTTGTGATGATGAAAGATATAGAAGACCATCTATTTTGGGTCAAACTG<br>TTCCATGTTCTACTACTACTATTAATGGTTCTACTCATGTTTTGAGAGA<br>TGCTGTTGAATGTGGTGGTCAATGGGAATCTCAATGTCATCCACCAGC<br>TACTTCTCCATTGGTTGCTGCTCAACCATCTGATACTTGA |
| gDAM6 | *yEGFP* | AAAACAATGAGCAAAGGAGAAGAACTTTTCACTGGAGTTGTCCCAATT<br>CTTGTTGAATTAGATGGTGATGTTAATGGGCACAAATTTTCTGTCAGA<br>GGAGAGGGTGAAGGTGATGCTACAATCGGAAAACTCACCCTTAAATTT<br>ATTTGCACTACTGGAAAACTACCTGTTCCATGGCCAACACTTGTCACT<br>ACTCTGACCTATGGTGTTCAATGCTTTTCCCGTTATCCGGATCACATG<br>AAAAGGCATGACTTTTTCAAGAGTGCCATGCCCGAAGGTTATGTACAG<br>GAACGCACTATATCTTTCAAAGATGACGGGAAATACAAGACGCGTGCT<br>GTAGTCAAGTTTGAAGGTGATACCCTTGTTAATCGTATCGAGTTAAAG<br>GGTACTGATTTTAAAGAAGATGGAAACATTCTCGGACACAAACTCGAG<br>TACAACTTTAACTCACACAATGTATACATCACGGCAGACAAACAAAAG<br>AATGGAATCAAAGCTAACTTCACAGTTCGCCACAACGTTGAAGATGGT<br>TCCGTTCAACTAGCAGACCATTATCAACAAAATACTCCAATTGGCGAT<br>GGCCCTGTCCTTTTACCAGACAACCATTACCTGTCGACACAAACTGTC<br>CTTTCGAAAGATCCCAACGAAAAGCGTGACCACATGGTCCTTCATGAG<br>TACGTAAATGCTGCTGGGATTACACATGGCATGGATGAGCTCTACAAA<br>TGA |

| | | |
|---|---|---|
| gDAM7 | *ADRB2* | AAAACAGAATTCAACGTTGGATCCAAGAATCAAAAATGTCTGATGCGG<br>CTCCTTCATTGAGCAATCTATTTTATGATGTGACGCAACAGAGAGACG<br>AGGTGTGGGTAGTCGGGATGGGTATCGTCATGAGCCTGATAGTTCTG<br>GCGATAGTCTTCGGAAACGTATTGGTGATTACAGCTATTGCGAAATTT<br>GAAAGACTACAAACCGTGACCAACTACTTCATTACTAGCCTAGCCTGT<br>GCTGATTTGGTAATGGGACTAGCGGTTGTACCGTTTGGCGCAGCGCA<br>TATTTTGATGAAGATGTGGACATTCGGTAATTTTTGGTGTGAGTTTTGG<br>ACATCTATTGATGTACTTTGCGTGACCGCCTCAATCGAAACATTGTGT<br>GTTATTGCGGTGGACCGTTATTTTGCCATTACCAGCCCATTTAAGTAT<br>CAGTCCCTGCTAACAAAGAATAAGGCACGTGTTATAATACTTATGGTA<br>TGGATTGTCTCTGGCCTAACCTCATTTCTGCCCATACAAATGCACTGG<br>TATCGTGCTACACATCAGGAGGCAATCAATTGTTACGCGAACGAGAC<br>GTGTTGCGACTTCTTCACAAACCAAGCGTATGCAATAGCTAGTTCCAT<br>AGTTTCATTTTACGTCCCTTTGGTAATAATGGTTTTTGTTTACTCCAGG<br>GTTTTCCAAGAAGCAAAGCGTCAGCTGCAAAAAATTGATAAATCAGAA<br>GGTCGTTTCCACGTTCAGAACCTTAGTCAAGTAGAGCAAGACGGTAG<br>GACTGGGCATGGCTTAAGAAGATCCTCCAAGTTCTGTCTGAAAGAACA<br>TAAGGCGTTGAAAACGCTGGGAATAATTATGGGCACATTTACTCTATG<br>TTGGTTGCCATTTTTCATCGTGAACATCGTCCATGTCATCCAAGACAAT<br>CTTATTAGAAAAGAAGTTTACATTCTACTGAACTGGATAGGGTATGTCA<br>ATTCCGGCTTCAATCCGCTTATCTATTGTAGATCCCCGGATTTCCGTA<br>TCGCTTTCCAAGAGCTACTGTGCCTACGTCGTAGTTCACTAAAAGCTT<br>ATGGCAATGGATACAGCTCAAACGGCAACACTGGTGAGCAATCAGGC<br>TATCACGTAGAGCAAGAAAAGGAGAACAAGTTATTATGTGAGGATCTT<br>CCGGGAACTGAAGACTTTGTCGGTCACCAGGGCACTGTCCCCTCTGA<br>TAACATTGATAGTCAGGGGGCGTAATTGTTCCACCAACGACTCTCTTCT<br>ATAA |

| | | |
|---|---|---|
| gDAM8 | *CXCR4* | AAAACAATGTCCATACCGTTGCCGCTTCTTCAGATATACACGTCTGAT<br>AATTACACGGAGGAGATGGGCTCTGGCGATTATGACTCTATGAAGGA<br>ACCCTGCTTTAGAGAAGAGAACGCAAACTTCAACAAAATTTTTCTGCC<br>TACGATATATTCCATAATATTCCTTACGGGGATCGTCGGCAATGGGTT<br>AGTTATACTGGTCATGGGTTACCAAAAGAAATTGCGTAGTATGACCGA<br>CAAATATCGTCTTCATTTGAGCGTTGCCGACTTGTTATTTGTCATCACA<br>CTGCCCTTTTGGGCTGTAGACGCCGTAGCGAACTGGTACTTTGGGAA<br>CTTTTTGTGTAAAGCTGTTCATGTTATCTACACCGTTAATCTATATTCCT<br>CCGTATTGATTTTAGCTTTTATATCCTTGGATAGATACCTGGCAATAGT<br>ACATGCAACGAATTCCCAACGTCCTAGAAAGTTACTGGCCGAGAAAGT<br>AGTGTACGTGGGGGTATGGATTCCGGCCCTGTTGTTGACGATACCGG<br>ATTTTATATTCGCTAATGTCAGCGAGGCAGACGATAGATATATATGCG<br>ACAGATTCTACCCTAACGACCTGTGGGTCGTTGTATTCCAATTCCAAC<br>ACATTATGGTGGGACTGATCTTGCCCGGCATAGTAATCTTGAGCTGCT<br>ATTGTATAATCATAAGTAAGCTGTCCCATAGTAAGGGACACCAAAAAA<br>GGAAAGCCTTGAAAACCACCGTGATACTAATTCTTGCTTTCTTTGCGT<br>GTTGGCTACCTTACTACATTGGGATAAGCATTGACTCTTTCATCCTGTT<br>AGAGATTATAAAGCAGGGATGTGAGTTCGAGAACACCGTGCACAAGT<br>GGATCTCCATCACTGAGGCGCTTGCATTCTTCCATTGTTGCCTTAACC<br>CCATTCTATATGCCTTCTTAGGCGCTAAATTCAAAACATCTGCTCAACA<br>TGCCCTTACTTCAGTTAGTAGAGGTAGTTCTCTGAAAATTTTGTCTAAG<br>GGTAAAAGGGGCGGACATTCCTCCGTTAGTACTGAAAGTGAATCATC<br>CTCTTTTCACAGCTCCTGA |

| | | |
|---|---|---|
| gDAM9 | *GLP-1R* | ATGGCCGGAGCACCCGGCCCCCTTAGGCTTGCTCTGCTGTTATTGGG<br>AATGGTTGGTAGGGCCGGACCAAGACCGCAAGGGGCCACGGTATCA<br>CTGTGGGAGACGGTTCAGAAGTGGAGAGAATATCGTAGGCAATGCCA<br>ACGTAGTCTAACAGAGGATCCGCCGCCGGCAACTGACCTTTTCTGCA<br>ACCGTACATTTGATGAATATGCTTGCTGGCCCGACGGCGAACCCGGA<br>AGCTTCGTAAATGTGTCATGTCCGTGGTACCTTCCATGGGCCTCTAGC<br>GTACCACAAGGGCATGTTTACAGATTCTGCACGGCCGAGGGATTATG<br>GTTACAAAAAGACAACTCCAGCTTACCCTGGCGTGATCTTTCTGAGTG<br>TGAGGAGTCTAAAAGAGGCGAACGTTCATCCCCCGAAGAGCAGTTGT<br>TGTTCTTGTACATAATTTATACGGTTGGTTATGCTCTTTCATTTTCCGC<br>ACTGGTGATCGCGTCAGCGATACTTCTGGGCTTTAGGCACTTACACT<br>GTACCAGGAACTACATACATCTGAATCTGTTCGCTTCTTTTATACTAAG<br>AGCTTTAAGTGTGTTCATTAAGGACGCCGCGCTAAAGTGGATGTATAG<br>CACTGCTGCTCAGCAGCACCAGTGGGACGGCTTGTTATCCTATCAAG<br>ACTCTTTATCATGCCGTTTGGTCTTTCTTCTGATGCAGTACTGTGTCGC<br>AGCCAATTACTATTGGCTTCTAGTCGAAGGGGTGTATCTGTACACATT<br>ACTAGCGTTTAGTGTCCTGTCTGAACAGTGGATTTTCAGATTATATGTA<br>TCTATCGGCTGGGGAGTCCCCCTACTTTTCGTTGTGCCATGGGGCAT<br>TGTAAAGTATCTGTACGAGGACGAGGGATGCTGGACAAGGAACTCAA<br>ACATGAATTATTGGCTGATTATCAGATTACCGATTCTATTCGCTATCGG<br>GGTTAATTTTCTAATTTTTGTAAGGGTGATTTGCATCGTTGTTAGCAAG<br>CTTAAAGCTAACCTTATGTGTAAAACCGACATAAAGTGCAGGTTGGCT<br>AAGTCAACTCTTACCCTTATCCCCCTACTGGGAACGCATGAAGTTATT<br>TTTGCTTTCGTCATGGATGAACACGCGCGTGGGACACTTCGTTTCATT<br>AAACTTTTCACGGAATTATCTTTTACCTCCTTTCAGGGGTTGATGGTAG<br>CGATTCTGTACTGCTTCGTTAACAATGAAGTACAGCTGGAATTCCGTA<br>AATCATGGGAGAGATGGAGACTGGAGCACCTGCACATACAACGTGAC<br>AGTTCAATGAAACCGTTGAAATGTCCCACTAGCAGTCTATCATCTGGA<br>GCCACCGCGGGCTCATCAATGTACACTGCGACCTGTCAGGCCTCTTG<br>CTCATGA |

| | | |
|---|---|---|
| gDAM10 | *CaSTE2* | ATGAACATTAATTCTACGTTCATACCAGACAAGCCTGGAGATATTATCA<br>TAAGTTACTCAATTCCCGGTTTGGACCAACCGATCCAAATCCCGTTTC<br>ATTCCCTAGACAGTTTTCAAACGGATCAAGCTAAGATCGCCCTAGTGA<br>TGGGCATAACGATAGGTAGTTGTTCCATGACACTTATCTTCCTGATTA<br>GCATTATGTACAAGACTAACAAATTAACGAACTTGAAGTTGAAATTAAA<br>GTTGAAGTACATTTTACAGTGGATCAATCAAAAGATCTTCACTAAGAAA<br>CGTAACGACAATAAGCAACAACAGCAGCAACAGCAACAGCAGATTGA<br>GTCAAGCAGTTATAATAATACGACAACTACGCTAGGCGGTTATAAGCT<br>TTTTCTTTTCTATCTTAACAGCCTGATACTGCTTATCGGGATAATTAGA<br>TCTGGGTGCTACTTAAATTATAACTTGGGACCATTGAACTCCCTTTCAT<br>TCGTTTTTACGGGATGGTATGATGGGAGTTCCTTTATATCTTCAGACG<br>TAACAAATGGTTTCAAGTGTATCCTATACGCTTTAGTCGAGATATCTTT<br>AGGATTTCAGGTATATGTGATGTTCAAAACGTCCAACTTGAAAATTTG<br>GGGCATAATGGCTTCCCTTCTGAGTATTGGACTAGGGCTTATCGTGGT<br>CGCATTCCAAATCAATTTGACTATCTTATCACATATCCGTTTTTCCAGA<br>GCTATAAGCACAAATAGAAGTGAGGAAGAATCCAGCTCATCTTTATCA<br>TCTGATTCCGTTGGATATGTCATAAATTCCATCTGGATGGACCTGCCA<br>ACGATTCTGTTTTCTATATCTATTAATATAATGACAATATTGCTAATTGG<br>TAAACTGATAATAGCAATCCGTACGAGAAGATACTTAGGATTGAAACA<br>ATTTGATTCTTTTCACATTCTTTTGATTGGGTTCTCACAAACTTTGATTA<br>TACCATCAATTATTCTAGTAGTCCATTATTTCTATCTATCTCAGAATAAG<br>GACTCTTTGTTACAACAAATTTCACTATTACTTATTATACTTATGTTGCC<br>CCTTTCAAGCTTGTGGGCGCAAACCGCTAACAATACACACAATATCAA<br>TAGTTCACCAAGCCTTTCCTTTATTAGCCGTCATCACTTATCCGACTCC<br>TCTCGTTCCGGCGGCTCCAACACTATCGTATCTAATGGCGGGTCTAAT<br>GGCGGCGGGGGAGGTGGTGGTAATTTTCCCGTCTCAGGCATAGATG<br>CCCAACTTCCTCCGGACATAGAGAAAATTCTGCATGAGGATAACAACT<br>ACAAGTTATTAAACTCCAATAATGAGAGTGTCAATGATGGAGACATAAT<br>TATTAACGATGAGGGCATGATAACAAAGCAGATTACAATCAAACGTGT<br>ATAG |

| | | |
|---|---|---|
| gDAM11 | *FgSTE2* | ATGTCCAAAGAAGTATTCGACCCATTCACGCAAAACGTCACTTTTTTTG<br>CACCGGATGGGAAGACAGAGATTTCTATCCCTGTTGCAGCTATTGAC<br>CAAGTTCGTAGGATGATGGTCAACACTACCATAAACTACGCAACGCAG<br>TTGGGGGCCTGTCTGATCATGCTTGTGGTGTTACTGGTCATGGTGCC<br>TAAGGAAAAGTTTAGGAGGCCGTTTATGATTCTTCAAATCACTAGTCTA<br>GTGATAAGCTGTTGTCGTATGTTACTACTGAGTATATTCCATTCCAGTC<br>AGTTTTTGGATTTCTACGTCTTCTGGGGAGACGACCATTCCCGTATCC<br>CTCGTAGCGCGTACGCACCAAGTGTCGCAGGCAACACGATGAGTTTG<br>TGCTTGGTAATCTCTGTCGAGACAATGCTAATGTCTCAAGCATGGACT<br>ATGGTTAGACTGTGGCCCAATGTTTGGAAATACATAATCGCGGGCGTA<br>AGTCTTATCGTTAGCATTATGGCTATTTCCGTGAGACTGGCGTACACG<br>ATTATACAAAACAATGCGGTATTGAAACTAGAGCCGGCATTTCATATG<br>TTCTGGCTAATAAAGTGGACTGTTATAATGAACGTTGCGAGCATTTCA<br>TGGTGGTGTGCAATATTCAATATCAAACTGGTTTGGCATCTGATATCTA<br>ATCGTGGAATTTTACCGTCATATAAAACATTTACACCAATGGAAGTTTT<br>AATAATGACAAACGGTATACTTATGATCATACCCGTGATCTTCGCCTC<br>ACTAGAGTGGGCCCATTTCGTAAACTTCGAGTCTGCCAGCCTAACCCT<br>TACTTCTGTGGCCGTCATACTGCCACTTGGAACCCTAGCCGCCCAGC<br>GTATCGCTAGTTCCGCTCCCTCTTCCGCAAACAGTACTGGCGCTAGTT<br>CAGGGATTCGTTACGGCGTGTCTGGTCCAAGTAGCTTCACAGGTTTC<br>AAAGCACCAAGTTTCAGCACGGGAACAACTGATAGACCCCACGTAAG<br>TATATACGCAAGGTGCGAGGCGGGTACGTCTTCCAGGGAACACATTA<br>ATCCTCAGGGAGTAGAACTAGCGAAACTGGACCCAGAGACAGACCAT<br>CACGTCAGAGTCGATAGAGCCTTTCTACAGAGAGAGGAGAGGATACG<br>TGCCCCCTTGTAG |

| | | |
|---|---|---|
| gDAM12 | *ZtSTE2* | ATGGTAGTTACCGCACCTCCCTCAGTCGACAGGACATACTTCATCCCT<br>AACAGCACCTTCGATCCATACCAGCAAGATTTGACATTGGTTTATCCC<br>GATGGAGTCCATGCATTGGTAGCAAATGTCGATGACATAGTGTACTTC<br>ATGGGATTAGCGGTCAAATCCACTCTGATATTTGCGATACAAATTGGT<br>ATATCTTTTGTTTTAATGTTGGTGATAGCATTACTGACCAAACCTGAAC<br>GTAGAGTGACCCTGGTCTTTTTCCTGAATATGACGGCCCTTTTTACTA<br>TTTTTATTAGGGCAATACTAATGTGTACAACATTCGTGGGCACCTACTA<br>TAATTTTTATAATTGGATTATGGGCAATTATCCAAACTCAGGTCTTGCT<br>GATCGTGTCTCTATCGCGGCTGAGGTCTTTGCCTTCTTAATTATTCTTT<br>CACTGGAGCTAAGCATGATGTTCCAGGTACGTATAGTTTGCATTAATC<br>TTAGTTCCTTCAGACGTAGAATTATCACATTTAGCAGTATTGTCGTCGC<br>GATGATAGTCTGTACTGTCAGGTTCGCCTTGATGGTGCTTTCCTGTGA<br>TTGGAGAATAGTAAATATAGGCGACGCAACCCAAGAGAAGAATCGTAT<br>TATCAATAGGGTAGCCAGCGGCTATAATATCTGTACCATCGCATCTAT<br>CATCTTCTTCAACACGATCTTCGTGAGCAAGTTGGCAGTGGCGATTAA<br>GCATAGAAGATCAATGGGGATGAAACAGTTTGGACCTATGCAGATTAT<br>TTTCGTAATGGGATGCCAGACACTGCTTATACCGGCTATCTTTGGCAT<br>TATCAGCTATTTCGCTCTTGCTAGCACGCAAGTTTACTCATTAATGCC<br>GATGGTGGTGGCAATCTTCTTGCCTCTTTCATCCATGTGGGCCAGTTT<br>CAATACTAACAAAACCAATTCTGTGACAAATATGAGGCAGCCCAACGT<br>CTACCGTCCTAACATGATAATAGGTCAGGACACGACGCAAAACAGCG<br>GTAAAAATACTAATATCTCAGGAACTAGTAACAGCACGGCTACCACCA<br>GCTCCTTTGCTTCTGATAAGAGGAGACTGAATCTAAGTTTTAATACCC<br>AGGGTACACTGGTCAACAGTATAAGTGAGGAAGAAGTCAACAACCCG<br>CAAAAATTGGGACCGAGTGCGACGGTAGCTGTTATGGATCGTGACAG<br>TCTAGAATTAGAGATGAGGCAACATGGCATAGCCCAAGGGCGTTCTT<br>ACTCAGTAAGGTCCGATTAG |

| | | |
|---|---|---|
| gDAM13 | *TrSTE2* | ATGGCCCCACACTTTGACCCCTTCCAACAAAGTGTTACCTTTCTGAGG<br>AGCGATGGTACAAGTTTTCCCATCTCCATGGCAGACTTCAACAAATTC<br>ATGCTGTACGCTGTGAGGACAAGCATCGCTGCAGCATCCCAGCTAGG<br>AGCGAGTGTTGTTATGGCTGTTCTATTGGCTCTATTGACCGCCCCAGA<br>TAAAAGACGTAGTATTGTGTTCTATTTGAATATTACCACTTTGCTAGTG<br>AACGTGTGTAGAACCCTGTCTACGACGATTTTCTTCACGTCATCATGG<br>GTTGAGATTTATACCTACTTTAGCGGGGACTATTCTCGTATAACAACG<br>GGCGCCTATGCCAATAGCGTAATGGGTACGATCGCTACGGGTATAAT<br>GGTTATTTTAATTGAGTTGTCACTGCTTATTCAAACGCACGTGTTGTGT<br>AGCACTCTGAGGGACCTATACAGAAATATTTTACTTGCTTGGTCTTGC<br>TTAGTTGCTGCGGTTCCTATCGCCTTTAGGATCGCATTCATGGTCGCT<br>AACGTTAAAGCGATTATGACGCAGTCTTCTCTGGGGAAGAACGTGTG<br>GATTCAGTCCAGTAGCAATATTTCCATTACAGTTTCCATATGCTACTTT<br>AGTTTGCTGTTCCTGGCAAAGTTGGGATATGCCATTTATACGAGGAGG<br>CTATTAGGTATGAAAGGCTTTGGAGTGATGCAGATCATTTTTATTATGG<br>CATGCCAAACGATGATCCTTCCCGCGATAATGTCCATCTTACAATATTT<br>CATACCGGAATTTGAGGTTAATACCAACATCTTAACCCTGTTGGCGTT<br>ATCTCTTCCGCTTACCACGCTTTGGTCCGCTGCGGCTGTGAGGCACG<br>GCCACAACAAGAACCAGGGTTCCGGCAGACATTTTTGGGGAGGTAGC<br>TCAGAGAAAAGTCTATTTGATCATAACAAAACCAGTGGGGGTAGCTTC<br>AGCCATCCCACTAGCACCTTAATAGGGTCTATGCCCGGACCCGAGAA<br>AGTCAAATCCGCCGACCACTTTGATAGACTGTACCCTGAACTACACGA<br>TACTGGAAATATCACGATTGAACGTAATTTCAGCGTCACTAGTAACAG<br>ATTG |

| | | |
|---|---|---|
| gDAM14 | *MsSTE3* | ATGTCAGGCTTTGCTACTCCGATCTTTGCTATACTGGCTGTTATCGTG<br>TCATTATTGCCGGTCCCGAGTCACTGGCGTGCGCGTAATTTCGTTATC<br>TTAGGGTTGGTTTTCTGGCTGGTTGTCGGTAACCTTAACATCTTTGTC<br>AACCGTATCATCTGGATGTCCAATGCCAAGAACTCCGCTCCAATATGG<br>TGTGACCTATCAGTTAAGTTGATGTCAATGGCCTCATTCTCTTTGCCG<br>TGCTCCGCGCTACTTATTTCTTCCAAACTTTATGATATAGCAAGCTTAA<br>AGTATTCCAAAAGGACCCCTGAAGAGAAGCAAAGGTTATGGATACTAG<br>AGCTGTTCTGCATTACGGTTTTGCCTGTTTTTTATTCACTGCTGACTTT<br>GGTCAGTCAGGGCCACAGGTTCAATATCGTATATGGCAGGGGTTGTG<br>AACCGGCCGTCTACTTTTCAAGTGTGTCCATCGTCATAGACTACGGTA<br>TCCCTGTATCTTTATGCATCACCTCCCTGGTTTATTCAGTTCTAAGCCT<br>GCGTCATTTTTTTATACATAAGAAAGACTTCGATGCTATTTTATCCAAG<br>AGTGGTTCTGGGATCAGCACAAAGAAGTTCCTAAGAATGATAAGCTTC<br>GCGTTAATCGACATACTTATAAATTTCCCAATACTGCTTGCAGCATTCG<br>CGCTAGAGGTGTCATACATGAAGATCATACCTTATACGTCCTGGGATT<br>TCGTTCATAAGAGGTTCTCAGACGTGTGGATTTACCCCAGGACTGCAA<br>TAAGCTCATCCAGATTTAAGCACTTTCTGACGATGACAAGCTTTGCAA<br>CGTGGTCTCAGTGTATGATGGGTTTTATATTTTTTTTCATGTTTGGATT<br>AAACACGGATATTAAGTCCGACTACGTAAAGGCCTTCATGAAGGTGAA<br>GGGGATGTTCCATCTTAAAACTCAGAAAGCAACACGTGAATACAAAGA<br>TGACAACATATGCTCAACTGATGACATTAGCGATGTGAGTAATCACAA<br>ATCAGAAGATTCTAACATCCACCCTGACGGAGTTTCAGTTGAGTTTAG<br>TCATGTTGATCTAGAAGCGAGTGGGCACGAAGCAAGTAATTATAGTAT<br>TTCAATTCCGAGCAACGATGACACGAAAACTAATCGTTTCGCT |
| gDAM15 | α-leader_P-factor | ATCTGTCATAAAACAATGAGATTTCCTTCAATTTTTACTGCTGTTTTATTCGCAGC<br>ATCCTCCGCATTAGCTGCTCCAGTCAACACTACAACAGAAGATGAAACGGCACA<br>AATTCCGGCTGAAGCTGTCATCGGTTACTCAGATTTAGAAGGGGATTTCGATGT<br>TGCTGTTTTGCCATTTTCCAACAGCACAAATAACGGGTTATTGTTTATAAATACTA<br>CTATTGCCAGCATTGCTGCTAAAGAAGAAGGGGTATCTCTCGATAAAAGAGAGG<br>CTGAAGCTACTTATGCCGATTTTCTGAGGGCATATCAGTCTTGGAACACGTTCGT<br>AAATCCAGATAGGCCCAATTTGTAAAGTGCAGGT |

| | | |
|---|---|---|
| gDAM16 | *mRuby2* | AAAACAATGGTGTCCAAAGGAGAGGAGTTAATCAAGGAAAACATGAGAATGAA<br>AGTTGTCATGGAGGGCTCCGTTAATGGTCACCAATTCAAGTGTACAGGGGAAG<br>GTGAAGGTAATCCTTACATGGGTACACAAACTATGAGAATTAAAGTAATTGAAG<br>GCGGACCACTACCATTTGCATTTGACATTCTGGCAACGTCATTCATGTACGGATC<br>ACGAACTTTCATCAAGTACCCTAAAGGTATACCAGACTTTTTCAAGCAATCTTTTC<br>CAGAGGGTTTTACATGGGAAAGGGTTACAAGATACGAAGATGGGGGTGTCGTC<br>ACAGTTATGCAAGATACTTCATTAGAAGATGGCTGCCTTGTCTATCATGTGCAAG<br>TAAGAGGGGTGAATTTTCCTTCTAACGGACCTGTGATGCAGAAAAAGACCAAAG<br>GTTGGGAACCAAATACTGAAATGATGTACCCAGCTGATGGAGGTTTGAGAGGC<br>TACACACACATGGCGCTTAAAGTTGATGGTGGAGGTCATTTGTCTTGTAGTTTTG<br>TTACCACTTATCGTTCTAAAAAGACTGTTGGCAATATCAAAATGCCAGGAATACA<br>TGCTGTAGACCACAGACTAGAAAGACTCGAAGAGAGCGATAACGAAATGTTCG<br>TTGTACAGAGAGAGCATGCCGTAGCCAAATTTGCTGGCTTAGGCGGTGGTATG<br>GATGAATTGTATAAGTAA |

| | | |
|---|---|---|
| DDgb009 | *gRNA_URA3_KO* | GGTTCATCATCTCATGGATCTGCACATGAACAAACACCAGAGTCAAAC<br>GACGTTGAAATTGAGGCTACTGCGCCAATTGATGACAATACAGACGAT<br>GATAACAAACCGAAGTTATCTGATGTAGAAAAGGATTAAAGATGCTAA<br>GAGATAGTGATGATATTTCATAAATAATGTAATTCTATATATGTTAATTA<br>CCTTTTTTGCGAGGCATATTTATGGTGAAGGATAAGTTTTGACCATCAA<br>AGAAGGTTAATGTGGCTGTGGTTTCAGGGTCCATAAAGCTTTTCAATT<br>CATCTTTTTTTTTTTGTTCTTTTTTTTTGATTCCGGTTTCTTTGAAATTTTT<br>TTGATTCGGTAATCTCCGAGCAGAAGGAAGAACGAAGGAAGGAGCAC<br>AGACTTAGATTGGTATATATACGCATATGTGGTGTTGAAGAAACATGA<br>AATTGCCCAGTATTCTTAACCCAACTGCACAGAACAAAAACCTGCAGG<br>AAACGAAGATAAATCAAAACTGTATTATAAGTAAATGCATGTATACTAA<br>ACTCACAAATTAGAGCTTCAATTTAATTATATCAGTTATTACCCGGGAA<br>TCTCGGTCGTAATGATTTTTATAATGACGAAAAAAAAAAAATTGGAAAG<br>AAAAAGCTTCATGGCCTTTATAAAAAGGAACTATCCAATACCTCGCCA<br>GAACCAAGTAACAGTATTTTACGGGGCACAAATCAAGAACAATAAGAC<br>AGGACTGTAAAGATGGACGCATTGAACTCCAAAGAACAACAAGAGTTC<br>CAAAAAGTAGTGGAACAAAAGCAAATGAAGGATTTCATGCGTTTGTAC<br>TCTAATCTGGTAGAAAGATGTTTCACAGACTGTGTCAATGACTTCACA<br>ACATCAAAGCTAACCAATAAGGAACAAACATGCATCATGAAGTGCTCA<br>GAAAAGTTCTTGAAGCATAGCGAACGTGTAGGGCAGCGTTTCCAAGA<br>ACAAAACGCTGCCTTGGGACAAGGCTTGGGCCG |

| | | |
|---|---|---|
| DDgb010 | *gRNA_HIS3_KO* | CTTCTCGACGTGGGCCTTTTTCTTGCCATATGGATCCGCTGCACGGTC<br>CTGTTCCCTAGCATGTACGTGAGCGTATTTCCTTTTAAACCACGACGC<br>TTTGTCTTCATTCAACGTTTCCCATTGTTTTTTTCTACTATTGCTTTGCT<br>GTGGGAAAAACTTATCGAAAGATGACGACTTTTTCTTAATTCTCGTTTT<br>AAGAGCTTGGTGAGCGCTAGGAGTCACTGCCAGGTATCGTTTGAACA<br>CGGCATTAGTCAGGGAAGTCATAACACAGTCCTTTCCCGCAATTTTCT<br>TTTTCTATTACTCTTGGCCTCCTCTAGTACACTCTATATTTTTTTATGCC<br>TCGGTAATGATTTTCATTTTTTTTTTTCCACCTAGCGGATGACTCTTTTTT<br>TTTCTTAGCGATTGGCATTATCACATAATGAATTATACATTATATAAAGT<br>AATGTGATTTCTTCGAAGAATATACTAAAAAATGAGCAGGCAAGATAAA<br>CGAAGGCAAAGTGACACCGATTATTTAAAGCTGCAGCATACGATATAT<br>ATACATGTGTATATATGTATACCTATGAATGTCAGTAAGTATGTATACG<br>AACAGTATGATACTGAAGATGACAAGGTAATGCATCATTCTATACGTG<br>TCATTCTGAACGAGGCGCGCTTTCCTTTTTTCTTTTTGCTTTTTCTTTTT<br>TTTTCTCTTGAACTCGAGAAAAAAAATATAAAAGAGATGGAGGAACGG<br>GAAAAAGTTAGTTGTGGTGATAGGTGGCAAGTGGTATTCCGTAAGAAC<br>AACAAGAAAAGCATTTCATATTATGGCTGAACTGAGCGAACAAGTGCA<br>AAATTTAAGCATCAACGACAACAACGAGAATGGTTATGTTCCTCCTCA<br>CTTAAGAGGAAAACCAAGAAGTGCCAGAAATAACAGTAGCAACTACAA<br>TAACAACAACGGCGGCTACAACGGTGGCCGTGGCGGTGGCAGCTTC<br>TTTAGCAACAACCGTCGTGGTGGTTACGGC |

| | | |
|---|---|---|
| DDgb008 | *pSNR52, GFP dropout, sgRNA, tSUP4* | tcggtctcagctgccaatgtctttgaaaagataatgtatgattatgctttcactcatatttatacagaaactt<br>gatgttttctttcgagtatatacaaggttattacatgtacgtttgaagtacaactctagattttgtagtgccct<br>cttgggctagcggtaaaggtgcgcatttttcacaccctacaatgttctgttcaaaagattttggtcaaacg<br>ctgtagaagtgaaagttggtgcgcatgtttcggcgttcgaaacttctccgcagtgaaagataaatgatctg<br>agaccgaaagtgaaacgtgatttcatgcgtcattttgaacattttgtaaatcttatttaataatgtgtgcgg<br>caattcacatttaatttatgaatgttttcttaacatcgcggcaactcaagaaacggcaggttcggatcttag<br>ctactagagaaagaggagaaatactagatgcgtaaaggcgaagagctgttcactggtgtcgtccctattc<br>tggtggaactggatggtgatgtcaacggtcataagttttccgtgcgtggcgagggtgaaggtgacgcaac<br>taatggtaaactgacgctgaagttcatctgtactactggtaaactgccggttccttggccgactctggtaac<br>gacgctgacttatggtgttcagtgctttgctcgttatccggaccatatgaagcagcatgacttcttcaagtcc<br>gccatgccggaaggctatgtgcaggaacgcacgatttcctttaaggatgacggcacgtacaaaacgcgt<br>gcggaagtgaaatttgaaggcgataccctggtaaaccgcattgagctgaaaggcattgactttaaagag<br>gacggcaatatcctgggccataagctggaatacaattttaacagccacaatgtttacatcaccgccgata<br>aacaaaaaaatggcattaaagcgaattttaaaattcgccacaacgtggaggatggcagcgtgcagctgg<br>ctgatcactaccagcaaaacactccaatcggtgatggtcctgttctgctgccagacaatcactatctgagc<br>acgcaaagcgttctgtctaaagatccgaacgagaaacgcgatcatatggttctgctggagttcgtaaccg<br>cagcgggcatcacgcatggtatggatgaactgtacaaatgaccaggcatcaaataaaacgaaaggctca<br>gtcgaaagactgggcctttcgttttatctgttgtttgtcggtgaacgctctctactagagtcacactggctca<br>ccttcgggtgggcctttctgcgtttataggtctcagttttagagctagaaatagcaagttaaaataaggcta<br>gtccgttatcaacttgaaaaagtggcaccgagtcggtgcttttgtttttatgtctggactgacaGCTGtg<br>agaccag |

### Supplementary Table S8

| Heterologous GPCR | Uniprot ID | Species of origin | Gα used in study | Amino acid sequence |
|---|---|---|---|---|
| *MTNR1A* | P48039 | Human | *GPA1* | MQGNGSALPNASQPVLRGDGARP<br>SWLASALACVLIFTIVVDILGNLLVIL<br>SVYRNKKLRNAGNIFVVSLAVADLV<br>VAIYPYPLVLMSIFNNGWNLGYLHC<br>QVSGFLMGLSVIGSIFNITGIAINRY<br>CYICHSLKYDKLYSSKNSLCYVLLI<br>WLLTLAAVLPNLRAGTLQYDPRIYS<br>CTFAQSVSSAYTIAVVVFHFLVPMII<br>VIFCYLRIWILVLQVRQRVKPDRKP<br>KLKPQDFRNFVTMFVVFVLFAICWA<br>PLNFIGLAVASDPASMVPRIPEWLF<br>VASYYMAYFNSCLNAIIYGLLNQNF<br>RKEYRRIIVSLCTARVFFVDSSNDV<br>ADRVKWKPSPLMTNNNVVKVDSV |
| *ADORA2B* | P29275 | Human | *GPA1* | MLLETQDALYVALELVIAALSVAGN<br>VLVCAAVGTANTLQTPTNYFLVSLA<br>AADVAVGLFAIPFAITISLGFCTDFY<br>GCLFLACFVLVLTQSSIFSLLAVAVD<br>RYLAICVPLRYKSLVTGTRARGVIA<br>VLWVLAFGIGLTPFLGWNSKDSAT<br>NNCTEPWDGTTNESCCLVKCLFEN<br>VVPMSYMVYFNFFGCVLPPLLIMLV<br>IYIKIFLVACRQLQRTELMDHSRTTL<br>QREIHAAKSLAMIVGIFALCWLPVH<br>AVNCVTLFQPAQGKNKPKWAMNM<br>AILLSHANSVVNPIVYAYRNRDFRY<br>TFHKIISRYLLCQADVKSGNGQAGV<br>QPALGVGL |

| | | | | |
|---|---|---|---|---|
| *HT4R* | Q13639-1 | Human | *GPA1/Gαi2 (DCGLF)* | MDKLDANVSSEEGFGSVEKVVLLT<br>FLSTVILMAILGNLLVMVAVCWDRQ<br>LRKIKTNYFIVSLAFADLLVSVLVMP<br>FGAIELVQDIWIYGEVFCLVRTSLDV<br>LLTTASIFHLCCISLDRYYAICCQPL<br>VYRNKMTPLRIALMLGGCWVIPTFI<br>SFLPIMQGWNNIGIIDLIEKRKFNQN<br>SNSTYCVFMVNKPYAITCSVVAFYI<br>PFLLMVLAYYRIYVTAKEHAHQIQM<br>LQRAGASSESRPQSADQHSTHRM<br>RTETKAAKTLCIIMGCFCLCWAPFF<br>VTNIVDPFIDYTVPGQVWTAFLWLG<br>YINSGLNPFLYAFLNKSFRRAFLIILC<br>CDDERYRRPSILGQTVPCSTTTING<br>STHVLRDAVECGGQWESQCHPPA<br>TSPLVAAQPSDT |
| *GLP-1R* | P43220 | Human | *GPA1/Gαs (QYELL)* | MAGAPGPLRLALLLLGMVGRAGPR<br>PQGATVSLWETVQKWREYRRQCQ<br>RSLTEDPPPATDLFCNRTFDEYAC<br>WPDGEPGSFVNVSCPWYLPWASS<br>VPQGHVYRFCTAEGLWLQKDNSSL<br>PWRDLSECEESKRGERSSPEEQLL<br>FLYIIYTVGYALSFSALVIASAILLGF<br>RHLHCTRNYIHLNLFASFILRALSVFI<br>KDAALKWMYSTAAQQHQWDGLLS<br>YQDSLSCRLVFLLMQYCVAANYYW<br>LLVEGVYLYTLLAFSVLSEQWIFRLY<br>VSIGWGVPLLFVVPWGIVKYLYEDE<br>GCWTRNSNMNYWLIIRLPILFAIGV<br>NFLIFVRVICIVVSKLKANLMCKTDIK<br>CRLAKSTLTLIPLLGTHEVIFAFVMD<br>EHARGTLRFIKLFTELSFTSFQGLM<br>VAILYCFVNNEVQLEFRKSWERWR<br>LEHLHIQRDSSMKPLKCPTSSLSSG<br>ATAGSSMYTATCQASCS |

| | | | | |
|---|---|---|---|---|
| *ADRB2* | P07550 | Human | *GPA1/Gαs (QYELL)* | MGQPGNGSAFLLAPNGSHAPDHDVTQ<br>ERDEVWVVGMGIVMSLIVLAIVFGNVL<br>VITAIAKFERLQTVTNYFITSLACADLVM<br>GLAVVPFGAAHILMKMWTFGNFWCEF<br>WTSIDVLCVTASIETLCVIAVDRYFAITSP<br>FKYQSLLTKNKARVIILMVWIVSGLTSFL<br>PIQMHWYRATHQEAINCYANETCCDFF<br>TNQAYAIASSIVSFYVPLVIMVFVYSRVF<br>QEAKRQLQKIDKSEGRFHVQNLSQVEQ<br>DGRTGHGLRRSSKFCLKEHKALKTLGIIM<br>GTFTLCWLPFFIVNIVHVIQDNLIRKEVYI<br>LLNWIGYVNSGFNPLIYCRSPDFRIAFQE<br>LLCLRRSSLKAYGNGYSSNGNTGEQSGY<br>HVEQEKENKLLCEDLPGTEDFVGHQGT<br>VPSDNIDSQGRNCSTNDSLL |
| *CXCR4* | P61073-1 | Human | *GPA1/Gαi2 (DCGLF)* | MEGISIYTSDNYTEEMGSGDYDSM<br>KEPCFREENANFNKIFLPTIYSIIFLT<br>GIVGNGLVILVMGYQKKLRSMTDKY<br>RLHLSVADLLFVITLPFWAVDAVAN<br>WYFGNFLCKAVHVIYTVNLYSSVLIL<br>AFISLDRYLAIVHATNSQRPRKLLAE<br>KVVYVGVWIPALLLTIPDFIFANVSE<br>ADDRYICDRFYPNDLWVVVFQFQH<br>IMVGLILPGIVILSCYCIIISKLSHSKG<br>HQKRKALKTTVILILAFFACWLPYYI<br>GISIDSFILLEIIKQGCEFENTVHKWI<br>SITEALAFFHCCLNPILYAFLGAKFK<br>TSAQHALTSVSRGSSLKILSKGKRG<br>GHSSVSTESESSSFHSS |

| | | | | |
|---|---|---|---|---|
| *MAM2* | Q00619 | *Schizosaccharomyces pombe* | *GPA1* | MRQPWWKDFTIPDASAIIHQNITIVSIVGEIEVPVSTIDAYERDRLLTGMTLSAQLALGVLTILMVCLLSSSEKRKHPVFVFNSASIVAMCLRAILNIVTICSNSYSILVNYGFILNMVHMYVHVFNILILLLAPVIIFTAEMSMMIQVRIICAHDRKTQRIMTVISACLTVLVLAFWITNMCQQIQYLLWLTPLSSKTIVGYSWPYFIAKILFAFSIIFHSGVFSYKLFRAILIRKKIGQFPFGPMQCILVISCQCLIVPATFTIIDSFIHTYDGFSSMTQCLLIISLPLSSLWASSTALKLQSMKTSSAQGETTEVSIRVDRTFDIKHTPSDDYSISDESETKKWT |
| *CaSTE2* | A0A1D8PTB4 | *Candida albicans* | *GPA1* | MNINSTFIPDKPGDIIISYSIPGLDQPIQIPFHSLDSFQTDQAKIALVMGITIGSCSMTLIFLISIMYKTNKLTNLKLKLKLKYILQWINQKIFTKKRNDNKQQQQQQQQQIESSSYNNTTTTTSGSYKLFLFYLNSLILLIGIIRSGCYLNYNLGPLNSLSFVFTGWYDGSSFISSDVTNGFKCILYALVEISLGFQVYVMFKTSNLKIWGIMASLLSIGLGLIVVAFQINLTILSHIRFSRAISTNRSEEESSSSLSSDSVGYVINSIWMDLPTILFSISINIMTILLIGKLIIAIRTRRYLGLKQFDSFHILLIGFSQTLIIPSIILVVHYFYLSQNKDSLLQQISLLLIILMLPLSSLWAQTANNTHNINSSPSLSFISRHHSFDSSRSGGSNTIVSNGGSNGGGGGGNFPVSGIDAQLPPDIEKILHEDNNYKLLNSNNESVNDGDIIINDEGMITKQITIKRV |

| | | | | |
|---|---|---|---|---|
| *FgSTE2* | I1RG07 | *Fusarium graminearum* | *GPA1* | MSKEVFDPFTQNVTFFAPDGKTEIS<br>IPVAAIDQVRRMMVNTTINYATQLG<br>ACLIMLVVLLVMVPKEKFRRPFMIL<br>QITSLVISCCRMLLLSIFHSSQFLDF<br>YVFWGDDHSRIPRSAYAPSVAGNT<br>MSLCLVISVETMLMSQAWTMVRLW<br>PNVWKYIIAGVSLIVSIMAISVRLAYT<br>IIQNNAVLKLEPAFHMFWLIKWTVIM<br>NVASISWWCAIFNIKLVWHLISNRGI<br>LPSYKTFTPMEVLIMTNGILMIIPVIF<br>ASLEWAHFVNFESASLTLTSVAVIL<br>PLGTLAAQRIASSAPSSANSTGASS<br>GIRYGVSGPSSFTGFKAPSFSTGTT<br>DRPHVSIYARCEAGTSSREHINPQ<br>GVELAKLDPETDHHVRVDRAFLQR<br>EERIRAPL |
| *ZtSTE2* | F9X131 | *Zymoseptoria tritici* | *GPA1, GPA1/Gα (MCGLI)* | MVVTAPPSVDRTYFIPNSTFDPYQQDLT<br>LVYPDGVHALVANVDDIVYFMGLAVKS<br>TLIFAIQIGISFVLMLVIALLTKPERRVTLV<br>FFLNMTALFTIFIRAILMCTTFVGTYYNFY<br>NWIMGNYPNSGLADRVSIAAEVFAFLIIL<br>SLELSMMFQVRIVCINLSSFRRRIITFSSIV<br>VAMIVCTVRFALMVLSCDWRIVNIGDA<br>TQEKNRIINRVASGYNICTIASIIFFNTIFV<br>SKLAVAIKHRRSMGMKQFGPMQIIFVM<br>GCQTLLIPAIFGIISYFALASTQVYSLMPM<br>VVAIFLPLSSMWASFNTNKTNSVTNMR<br>QPNVYRPNMIIGQDTTQNSGKNTNISG<br>TSNSTATTSSFASDKRRLNLSFNTQGTLV<br>NSISEEEVNNPQKLGPSATVAVMDRDSL<br>ELEMRQHGIAQGRSYSVRSD |

| | | | | |
|---|---|---|---|---|
| *TrSTE2* | A0A022VRI2 | *Trichophyton rubrum* | *GPA1, GPA1/Gα (LCGLI), GPA1/Gα (DSGIL)* | MAPHFDPFQQSVTFLRSDGTSFPISMA<br>DFNKFMLYAVRTSIAAASQLGASVVMA<br>VLLALLTAPDKRRSIVFYLNITTLLVNVCR<br>TLSTTIFFTSSWVEIYTYFSGDYSRITTGAY<br>ANSVMGTIATGIMVILIELSLLIQTHVLCS<br>TLRDLYRNILLAWSCLVAAVPIAFRIAFM<br>VANVKAIMTQSSLGKNVWIQSSSNISIT<br>VSICYFSLLFLAKLGYAIYTRRLLGMKGFG<br>VMQIIFIMACQTMILPAIMSILQYFIPEFE<br>VNTNILTLLALSLPLTTLWSAAAVRHGH<br>NKNQGSGRHFWGGSSEKSLFDHNKTS<br>GGSFSHPTSTLIGSMPGPEKVKSADHFD<br>RLYPELHDTGNITIERNFSVTSNRL |
| *MsSTE3* | A0A1M8A1X3 | *Malassezia sympodialis* | *GPA1, GPA1/Gα (ETGFL)* | MSGFATPIFAILAVIVSLLPVPSHWRARN<br>FVILGLVFWLVVGNLNIFVNRIIWMSNA<br>KNSAPIWCDLSVKLMSMASFSLPCSALLI<br>SSKLYDIASLKYSKRTPEEKQRLWILELFCI<br>TVLPVFYSLLTLVSQGHRFNIVYGRGCEP<br>AVYFSSVSIVIDYGIPVSLCITSLVYSVLSLR<br>HFFIHKKDFDAILSKSGSGISTKKFLRMIS<br>FALIDILINFPILLAAFALEVSYMKIIPYTS<br>WDFVHKRFSDVWIYPRTAISSSRFKHFL<br>TMTSFATWSQCMMGFIFFFMFGLNTDI<br>KSDYVKAFMKVKGMFHLKTQKATREYK<br>DDNICSTDDISDVSNHKSEDSNIHPDGV<br>SVEFSHVDLEASGHEASNYSISIPSNDDT<br>KTNRFA |
